# Testing a theoretical framework for the environment-species abundance paradigm: a new approach to the Abundant Centre Hypothesis

**DOI:** 10.1101/2022.01.03.474819

**Authors:** Alakananda Maitra, Rohan Pandit, Mansi Mungee, Ramana Athreya

## Abstract

The linkage between environment, a species’ fitness and its abundance is central to the theory of evolution. So far, all studies of this linkage have been heuristic and empirical due to an inability to determine fitness either experimentally (independent of abundance) or theoretically (from species-environment interaction). One category of such studies involves the Abundant Centre Hypothesis which posits that a species’ abundance rises to a maximum at the centre of its range. We argue that the confusing mix of results from ACH studies arises from ignoring the central premise that the abundance distribution cannot be independent of the environment. First, we employed a theoretical framework to identify an environmental context (an elevational transect; 200-2800 m in the eastern Himalayas) likely to favour ACH. We then improved upon some previously identified conceptual and methodological shortcomings of ACH studies. Using systematically collected bird data (245 species; 15867 records) from that transect we found that the community average abundance profile is symmetric, as expected by ACH. Notwithstanding which, the abundance profiles of individual species showed a small degree of asymmetry which was correlated with elevation. This elevational dependence may be due to the hard elevational limits at the lower and upper ends of the mountain, as expected from theoretical considerations. We also showed that the average abundance profile shape is close to gaussian, while ruling out uniform and inverted-quadratic shapes. This work demonstrates that selecting a particular category of environmental contexts can help in integrating theoretical tools into a field dominated by empirical studies. Such a union should spur the development of more detailed and testable theoretical models for better insights in an important field.

## Introduction

The Environment-Traits-Fitness-Abundance linkage is a central tenet of the theory of evolution. Yet, complex geographical pattern of environmental parameters, intricate interplay of multiple traits of a species, and our inability to “view the environment through a species’ eyes” make it difficult, if not impossible, to translate an observed environment-trait(s) pattern into a fitness profile for most species. Furthermore, fitness cannot be inferred independently of abundance. Species abundance studies have remained entirely heuristic and empirical because of this inaccessibility of fitness from both theoretical and observational sides. The several hundred environment-abundance studies till date (of Abundant Centre Hypothesis and Niche modeling) have only yielded a confusing welter of results. We suggest that the absence of theoretical inputs, due to the complicated and intractable differential equations needed to model a typical, complex two-dimensional environmental gradient, has been a key factor for the lack of progress. Here, we show that inputs from tractable theoretical models can be utilised if appropriate environmental gradients are chosen for study.

Abundant Centre Hypothesis (ACH), that *the abundance peak of a species coincides with the centre of its distribution*, is the most commonly tested macro-ecological pattern of the environment-abundance paradigm (Brown, 1984; Sagarin & Gaines, 2002a; Murphy et al., 2006; Rivadeneira et al., 2010; Fenberg & Rivadeneira, 2011; Baldanzi et al., 2013; Freeman, 2017; Pironon et al., 2017; Burner et al., 2019; Wen et al., 2020). Niche modeling went further to determine correlations between observed abundance and (a large number of) environmental variables (e.g. VanDerWal et al., 2009; Martínez-Meyer et al., 2012; Dallas et al., 2020). However, both kinds of studies have been almost entirely heuristic and empirical in nature.

An earlier review found support for ACH in only 39% of 145 direct tests conducted in 22 field studies (Sagarin & Gaines, 2002b); the situation has not improved in the last two decades. In an excellent review, Santini et al., 2019 identified a number of issues with the way ACH has been studied so far. These include confounding the geographic/geometric and environmental/ecological definitions of a species range, multiple climate variables with two-dimensional gradient (also Sagarin & Gaines, 2002a), confusion in terminology and definitions (also Borregaard & Rahbek, 2010), data quality (heterogeneity and insufficient normalisation for effort and species ecology), incomplete sampling of species ranges, and difficulty in separating location-specific patterns from peculiarities of particular species (Borregaard & Rahbek, 2010).

Many of the issues listed are due to logistical and/or resource constraints. However, we have identified two core issues which arise from the entirely heuristic and empirical approaches employed so far: (i) None of the previous studies have asked if ACH should at all have been expected at their sites, (ii) Confusion in identifying the “centre” of a species distribution, largely due to the conflation of geography with environment. Figure 1 shows the global distribution of a species (from our study). Does the geometric centroid of the distribution have any relevance when it may not even fall within it? However, the absence of a theory precludes the calculation of an ecological centroid. How will patchiness of occurrence within the distribution, possibly at mutliple scales, change the analysis and conclusion? Weighting the locations within the distribution by the abundance to determine its centroid creates logical circularity (Sagarin et al., 2006). Santini et al., 2019 weighted the locations using inputs from niche models (which itself is entirely empirical) but that did not improve the conclusions. We concur with previous researchers that the only secure conclusion from the results till date is that A*CH cannot be valid for all environment-species contexts* (e.g. Sagarin et al., 2006; Gaston, 2009).

**Figure 1:**
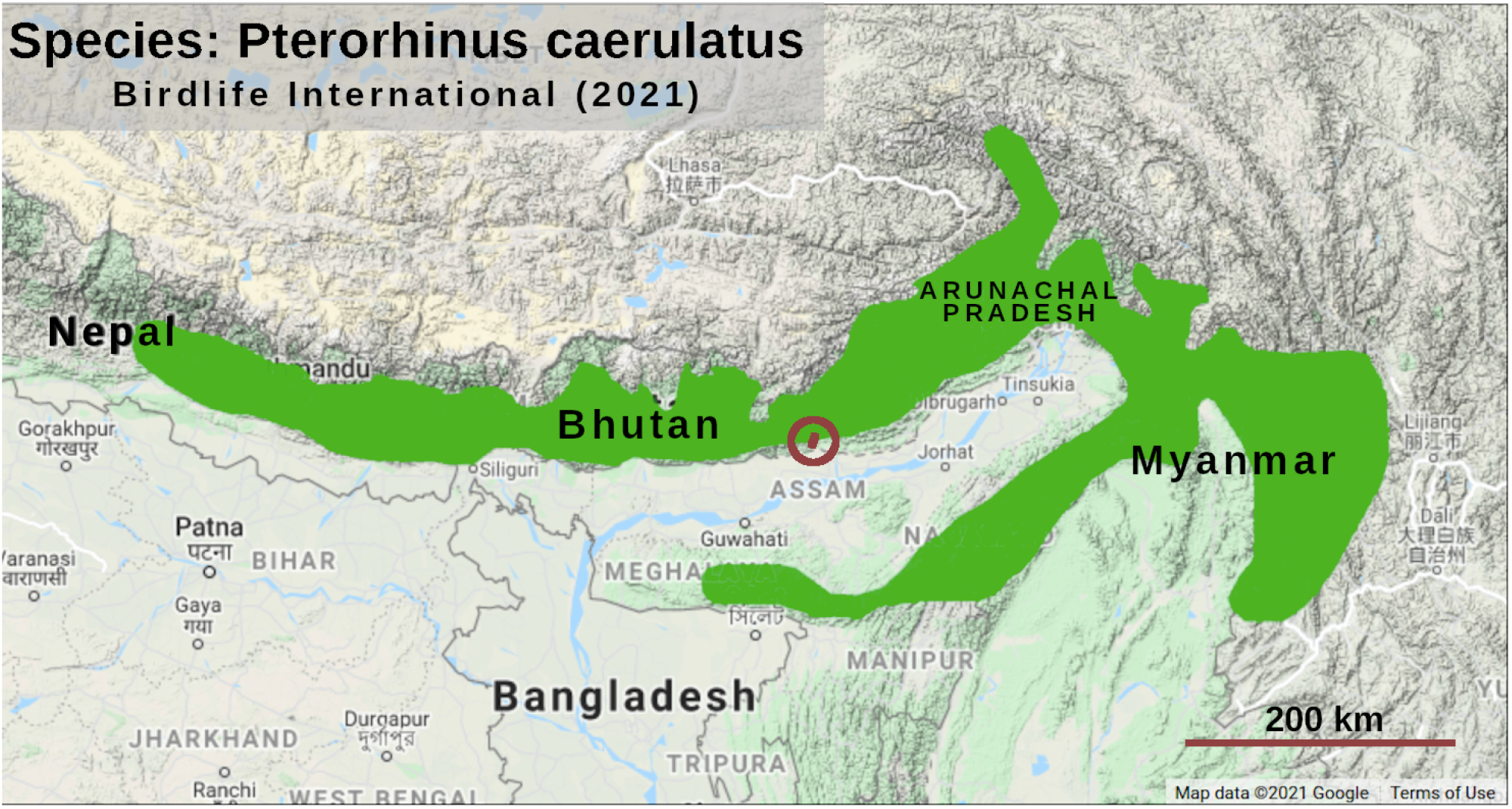
The global distribution of a bird species (bright green). The geometric centroid of the complex shape has no ecological relevance as it may even lie outside the distribution. Pockets of absence of this species within the envelope will further complicate the issue. The sampled elevational transect lies within the spot inside the circle to the right of Bhutan.

### Our approach differs from previous ones in three ways

First, we used an available theoretical framework (Kirkpatrick & Barton, 1997; hereafter KB97) to identify an environmental context (a steep elevational transect in a large mountain chain) for which ACH emerged as a prediction. Of course, every environment-species context will have an associated abundance distribution as a testable prediction. However, dealing with the symmetry implicit in ACH has observational and theoretical advantages. We also suspect that, generally, predictions of ACH may be associated with simpler and hence mathematically more tractable environmental contexts. This would allow theory and observations to progress together and sustain each other. Even the rejection of ACH by data could contribute to progress by identifying inappropriate assumptions in the model.

Then, we collected through field observations a large amount of primary abundance data (245 bird species; 15867 individuals) across a large environmental gradient (2600 m elevational transect in the eastern Himalayas) in a systematic manner (47 equispaced elevations under similar habitat visibility, 24 replicates matched for time of day across elevations; inside 3 years; by the same observer for uniformity).

Finally, we shifted the reference location for characterising range parameters from the periphery to the abundance peak. Range edges are associated with small, fluctuating, sink populations which are statistically unreliable (Hengelveld & Haeck, 1982; Brown, 1984; Lawton, 1993; Hoffmann & Blows, 1994), while the large number of records at the abundance peak makes its location statistically more stable.

### We used the following components to address some of the afore-mentioned issues

1. Theoretical framework: We started with KB97 to link an environmental gradient to a particular abundance profile using standard ecological processes and the life-history traits of the species. The formalism is applicable to a one-variable (univariate) environment with a gradient along one geographical dimension. Multiple environmental variables can effectively univariate if they are strongly correlated.
2. Study site: Effectively, our study site along an elevational transect had a one-dimensional environmental gradient and was univariate (with elevation being the predictor for multiple environmental variables). Its compact size (projected rectangle 6 km x 15 km) avoided the impact of confounding variables like zoo-geographical history, geographical climate variability, etc. The elevational transect spanned the entire *local* environmental range of many dozens of species.
3. Symmetry: KB97 predicted ACH (under certain assumptions) for the linear environmental gradient at the site.
4. A large and systematically collected abundance data set, as mentioned earlier.
5. Modified metric for ACH: Instead of coincidence between the geometric midpoint (of the outermost records) and the abundance peak, we tested ACH by the symmetry of half-range widths on either side of the abundance peak (Figure 2). This metric is more robust because it shifts the reference location from the sparsest regions to the densest. Secondly, the two half-range widths (quantified in several ways) were estimated using all the data rather than the distance to just the two farthest records.
6. Multiple taxa: We targeted the entire bird community in a species-rich eastern Himalayan site in the expectation that the average over all species should cancel the asymmetries introduced into individual profiles by competitive interactions between species pairs, and so reflect the impact of the environment.

**Figure 2:**
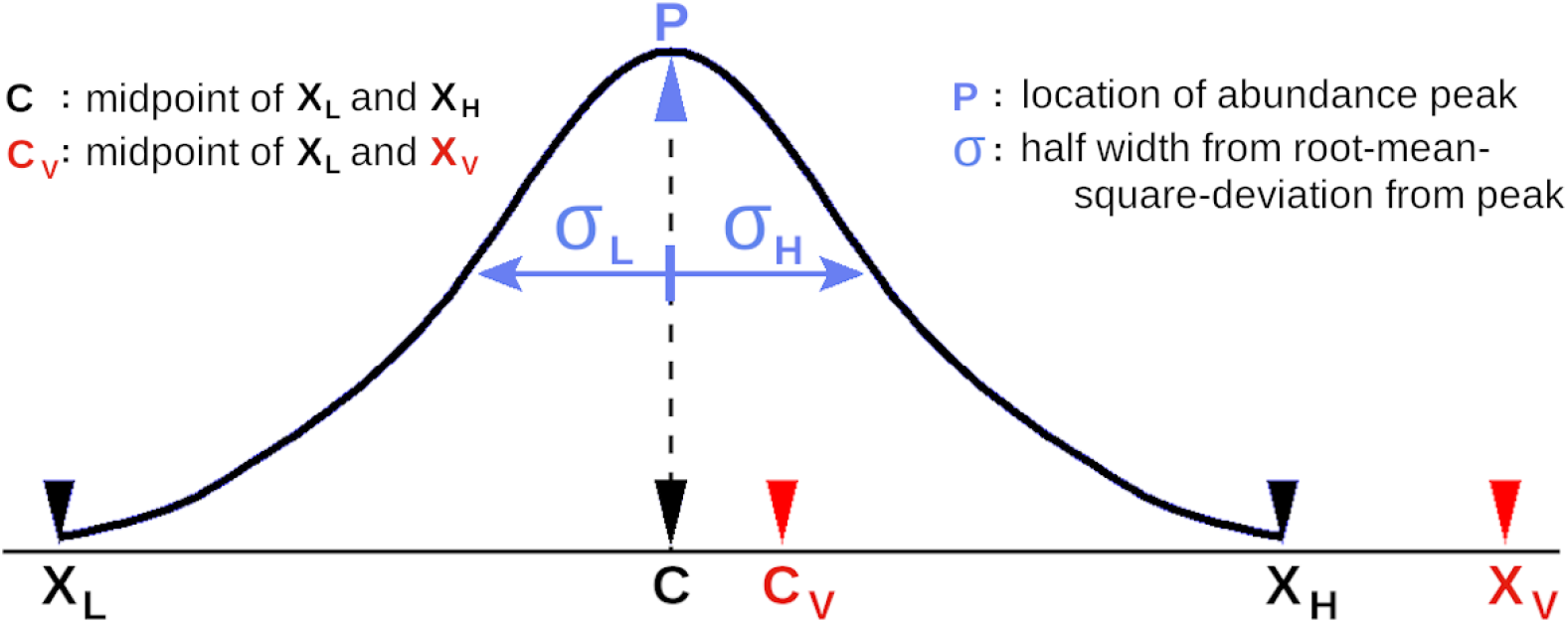
Metrics from the species abundance profile. X_L_ and X_H_ are the outermost records for a “good” distribution. **ACH so far**: the centre is expected to coincide with the peak. The centre is determined from the two outermost records, ignoring all other records. However, even a single vagrant (X_V_) can change the centre substantially (to C_V_). **ACH used in this work**: the half widths (σ_L_ and σ_H_) on either side of the peak are calculated using all the data. ACH is equivalent to σ_L_ = σ_H_. The few peripheral points have little impact on the estimates of the peak and the half-widths.

KB97 proposed a one-dimensional theoretical framework for the environment-abundance paradigm by incorporating several ecological processes like genetic diversity, directional selection of traits, vagility, etc, into the heat diffusion equation. Solving the equation under different sets of assumptions yields different abundance profiles. It predicts a gaussian^1^ abundance profile for (i) linearly increasing trait discrepancy (the difference between the traits of the local population and environmental optimum) along the environmental gradient (ii) quadratic relationship between fitness and trait discrepancy, and (iii) exponential relationship between abundance and fitness. Steep elevational transects may be expected to result in linear trait discrepancies. So, we tested two linked predictions of KB97 using records of all the birds encountered along an elevational transect on a single mountain in the eastern Himalayas.

We tested the two predictions separately since they are related to two different assumptions in KB97:

i. abundance profiles are symmetric about the abundance peak, i.e. ACH.
ii. abundance profiles are gaussian (i.e. have a peak and long tails); and not ∩-quadratic (a peak but no tails) or uniform (neither peak nor tail).

If the data were to reject ACH either (i) one or more of the assumptions listed above are inappropriate or (ii) KB97 needs to incorporate one or more relevant ecological process.

However, testing ACH is a secondary objective, and only a tool to demonstrate that there are environmental contexts which can accommodate theoretical tools. Indeed, one should move on from *ACH-or-not* question to understand profile shapes along different environmental gradients.

## Methods

See supplementary material for more details on each of the paragraphs below

We recorded bird abundance along a compact transect (elevation 200-2800 m asl, projected area 15 km x 6 km) in Eaglenest wildlife sanctuary in the eastern Himalayas, Arunachal Pradesh, north-east India (Athreya, 2006; Mungee, 2018). Eaglenest hosts contiguous pristine habitats ranging from tropical-evergreen at 100 m elevation to temperate forests at 3250 m. A vehicle track provides access to 100-2780 m on the southern slope of the mountain.

The same observer recorded all the birds encountered in 200 m line transects (along the road) at 47 elevations between 500m and 2800m, at elevational intervals of 50 m (Figure 3; Suppl. Table ST1). Each transect was sampled during a 5+5 minute traverse along the road, on 12 different days between 2^nd^ May and 3^rd^ July, 2012-2014 (Suppl. Figure S1). All individuals detected (visually and aurally) within 20 m from the path were recorded. Due to logistical issues at elevations below 500 m in Eaglenest, we sampled 4 transects (12 replicates each) at 200 m elevation in neighbouring Pakke Tiger Reserve, 25 km away. The 200 m data was only used to determine if a species distribution extended below 500 m.

**Figure 3:**
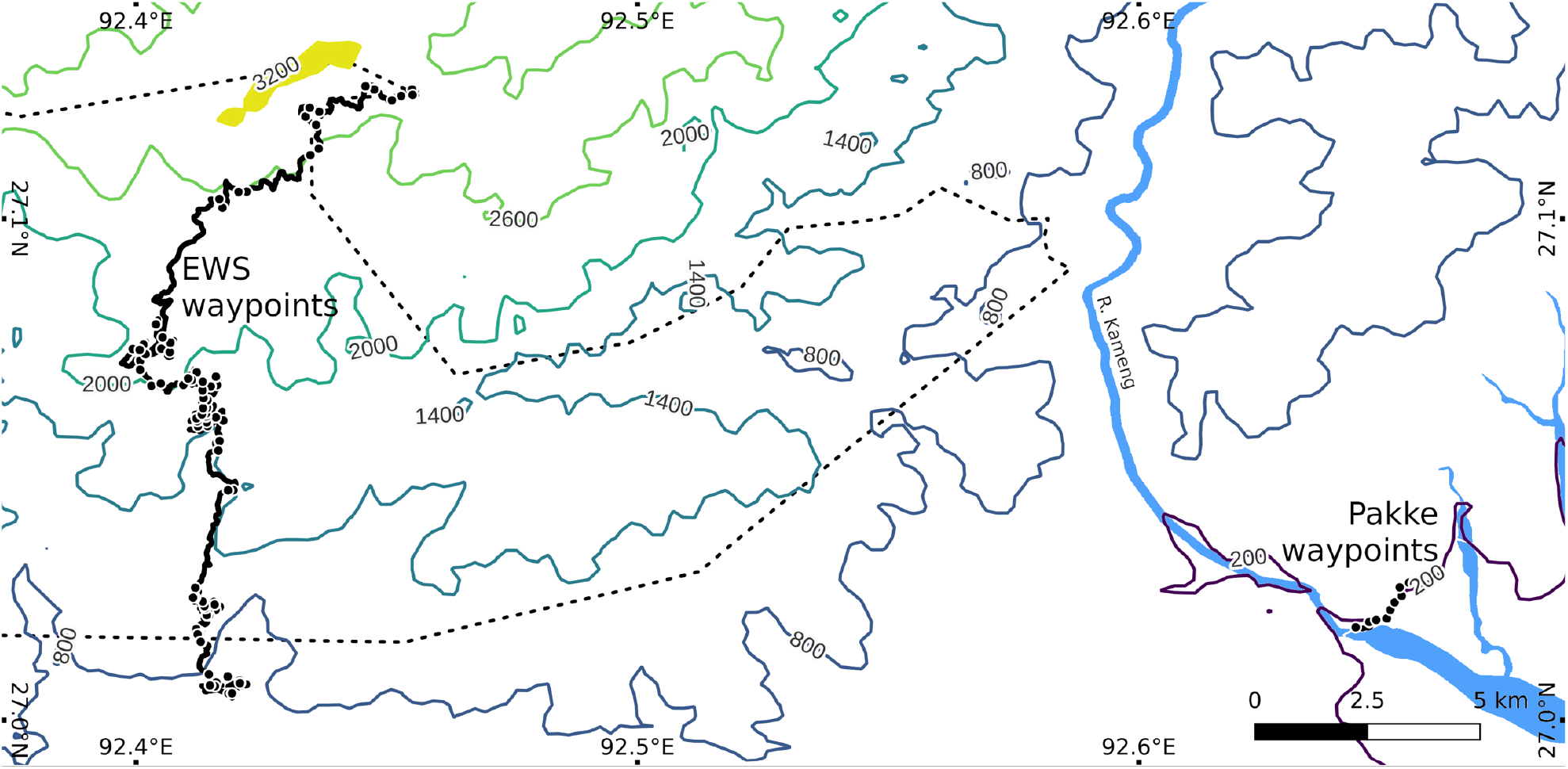
Study site in Eaglenest wildlife sanctuary, Arunachal Pradesh, India. The plot shows the elevational contours and sampling locations. The 49 Eaglenest transects in 500-2800 m are along a vehicle track. The Eaglenest ridge at 3200-3250 m is shown in yellow.

### Statistical Parameters of Abundance Profiles

The abundance peak elevation was located using a cubic fit to the smoothed profile (Suppl. Figure S2). We calculated the root-mean-square-deviation from the peak on the lower (σ_LS_) and upper (σ_HS_) sides to define asymmetry as

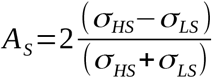

A_S_ is zero for symmetric profiles and ranges between –2 and +2. A_S_ has a simple relationship with prevalent mathematical definitions of skewness while being less error-prone for our particular case (Suppl.Figure S3). For a bigaussian profile (Wallis et al., 2014; Suppl. Figure S2) – the asymmetric counterpart of the gaussian – an identical value of A can be obtained by substituting σ_LS_ and σ_HS_ by other parameters like (i) abundance on either side of the peak, N_L_ and N_H_, for the number asymmetry A_N_, and (ii) the distances over which abundance declines to 60.65% of the peak, σ_L60_ and σ_H60_, for the asymmetry A_60_ in the inner regions of the distribution. Estimates of A_S_ and A_N_ are impacted by any section of the profile extending beyond the sampled range. A_S_ is sensitive to the distance of the records from the peak while A_N_ is not.

We calculated the Pearson and Spearman’s correlation coefficient for asymmetry and modal elevation, and also the linear regression between them.

We estimated the errors of all profile, regression and correlation parameters by simulating 400 profiles for each species and processing them in the same manner as the observed data. The Monte Carlo simulations used the smoothed observed distribution as the model, poisson overdispersion factor 2.0 (Suppl. Figure S4) and a negative binomial random number generator (Lindén & Mäntyniemi, 2011; *rnbinom in R*; R Core Team, 2020). The Poisson overdispersion factor also takes care of the detectability of birds.

We quantified profile shapes using kurtosis (K) which is characteristic of each family of curves independent of their mean and SD: K_G_ = 3.0 for gaussian, K_Q_= 2.14 for ∩-quadratic, and K_U_ = 1.8 for uniform profiles. Kurtoses, which depend on the 4^th^ power of the coordinate, typically have large errorbars. Therefore, we calculated kurtosis for the species-averaged community profiles in 3 elevational bands (800-1450 m, 1451-1820 m, and 1821-2400 m). The expected values for the smoothed and species-averaged community profiles are K_CG_ = 3.0 (gaussian), K_CQ_ = 2.23 and K_CU_ = 1.93 (uniform).

Only species which statisfied the following criteria were included in the analysis:

i. Total abundance ≥ 30
ii. Number of elevations with non-zero records ≥ 5
iii. Profile shape: unimodal when smoothed with full-width up to 1.5 x SD
iv. The level of the smoothed abundance profile at the sampling edge is:

- less than 5% of the peak – not truncated: A_S_, A_N_, A_60_, and kurtosis
- between 60% and 5% of the peak - partially truncated: only A_60_

All analyses were done using scripts written for the computing platform R (R Core Team, 2020).

## Results

Of the 245 species (15867 individuals) recorded, 44 satisfied the criteria for all 3 asymmetry metrics, and we calculated only A_60_ for another 19 species. Example profiles are shown in Suppl. Figure S5.

The statistics of the community averaged asymmetry metrics are shown in Table 1. The mean asymmetries varied between 1.6% and 9.9% for the three metrics but symmetry cannot be ruled at the 95% confidence level in any of them. The 95% confidence interval of only 5 out of 63 species did not include A = 0. This is consistent with the expectation of 3±3.5 outliers purely from stochasticity.

**Table 1.** Community mean asymmetry of the abundance profiles for birds in Eaglenest, Eastern Himalayas, India. The community mean asymmetry is consistent with the value zero, i.e. supports ACH. The SD of error-normalised asymmetry suggest that, compared to a gaussian, the average profile is more sharply peaked near the centre (A_60_ < 1.0) and has a heavier tail (A_S_ > 1.0) – i.e. leptokurtic.

| Asymmetry | N | Mean | SE | CI 95% | Error-normalised Asymmetry | SD |
| --- | --- | --- | --- | --- | --- | --- |
| $A_S$ | 44 | 0.073 | 0.117 | (-0.047, 0.356) | $Z_S$ | 1.29 |
| $A_N$ | 44 | -0.016 | 0.087 | (-0.157, 0.243) | $Z_N$ | 0.87 |
| $A_{60}$ | 63 | 0.099 | 0.051 | (-0.141, 0.156) | $Z_{60}$ | 0.80 |

The distributions of error-normalised asymmetry metrics z_**i**_ = A_**i**_/ε_**i**_, where A_i_ is the asymmetry for the i-th species and ε_**i**_ its error estimate, are shown in Figure 4 (also Table 1). Most of the values lie betweeen ±2 which suggests that the dispersion seen in the plot is consistent with that due to measurement errors. The SD was expected to be ~1.0 for dispersion dominated by measurement errors, which is approximately the case. The SDs are 0.80 for A_60_ (only data from near the abundance peak) and 1.29 for A_S_ (all data).

**Figure 4.**
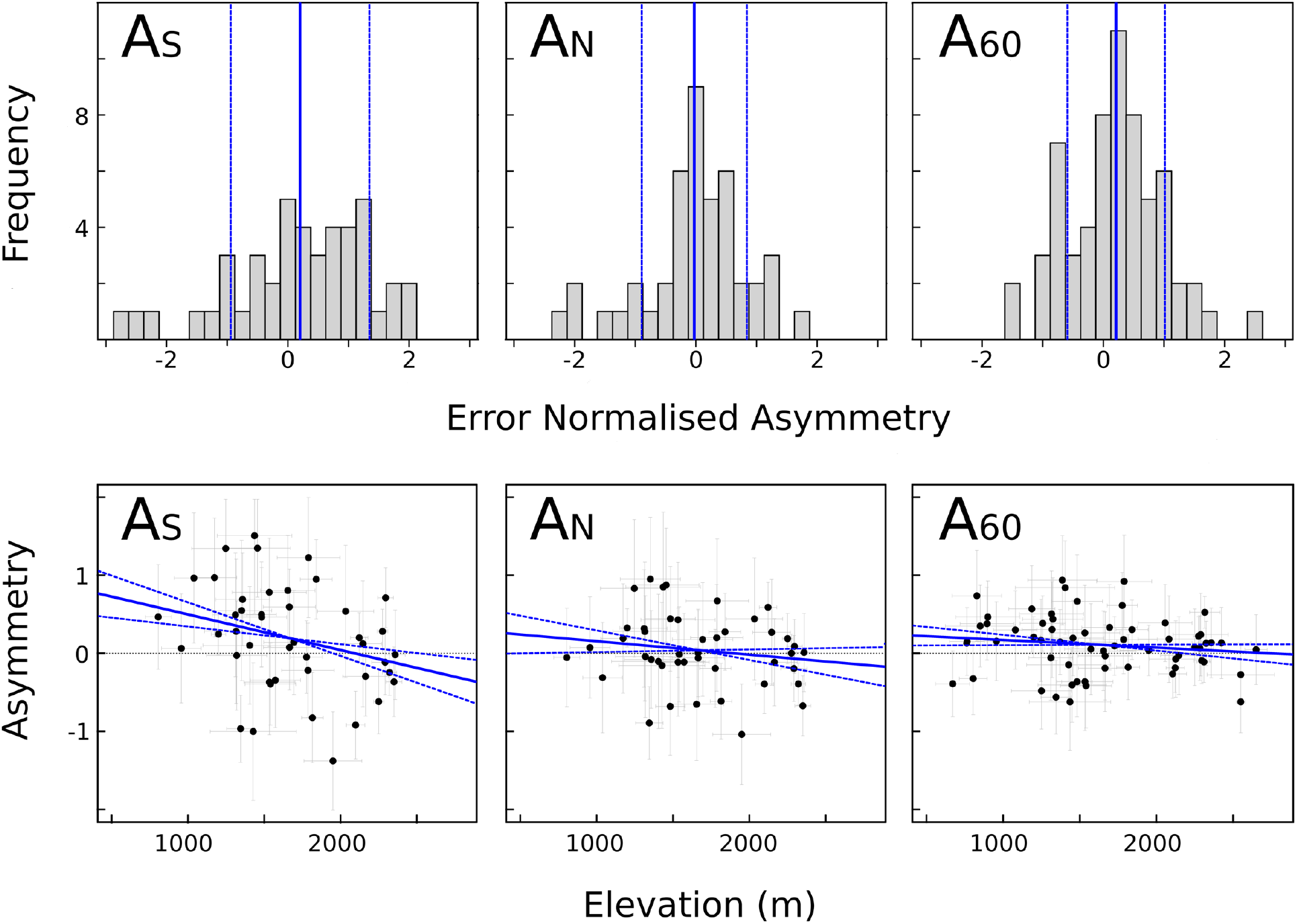
Species abundance profile asymmetry. A_S_: RMSD-based asymmetry, A_N_: count-based asymmetry, A_60_: Scale-length-based asymmetry. **Upper row**: Histograms of error-normalised asymmetry for each species (z = A/ε, A: asymmetry and ε: its error estimate). The vertical blue lines represent the mean (solid) and ±1 SD (dashed). The statistics of the distributions are in Table 1. **Lower row**: Linear regression between asymmetry and modal elevation. The blue lines are the best fit (solid) and ±1 SE (dashed) models. The regression parameters are in Table 2.

The correlation coefficient and linear regression for the asymmetry-elevation relationship are shown in Table 2 and Figure 4. Both correlation and regression analysis show a significant dependence of A_S_ on elevation at the 95% confidence level, but not for A_N_ and A_60_.

**Table 2.** Relationship between elevation and Asymmetry. The 95% confidence intervals were determined by Monte Carlo simulations of the observed profiles

| Variables | N | Pearson Correlation |  | Spearman Rank Correlation |  | Linear Regression |  |  |
| --- | --- | --- | --- | --- | --- | --- | --- | --- |
|  |  | r | CI 95% | r | CI 95% | Slope | Intercept | R<sup>2</sup> |
| $A_S \sim E_M$ | 44 | -0.321 | (-0.52, -0.08) | -0.34 | (-0.54, -0.10) | $(-5.91 \pm 2.34) \times 10^{-4}$ | $1.127 \pm 0.43$ | 0.085 |
| $A_N \sim E_M$ | 44 | -0.179 | (-0.38, 0.06) | -0.197 | (-0.40, 0.02) | $(-1.76 \pm 2.13) \times 10^{-4}$ | $0.33 \pm 0.40$ | 0.024 |
| $A_{60} \sim E_M$ | 63 | -0.153 | (-0.37, 0.10) | -0.198 | (-0.43, 0.06) | $(-1.03 \pm 1.04) \times 10^{-4}$ | $0.28 \pm 0.19$ | 0.018 |

The average community profiles in the 3 elevational bands are shown in Figure 5a. The kurtoses of their half profiles (on either side of the peak) are plotted in Figure 5b. The mean kurtosis for the six half-profiles was 3.81 (CI_95_ [2.85, 4.76]), which is consistent with a gaussian profile (K_CG_ = 3.0). These measurements rejected ∩-quadratic (K_CG_ = 2.23) and uniform (K_CU_ = 1.93) distributions with p < 0.01.

**Figure 5.**
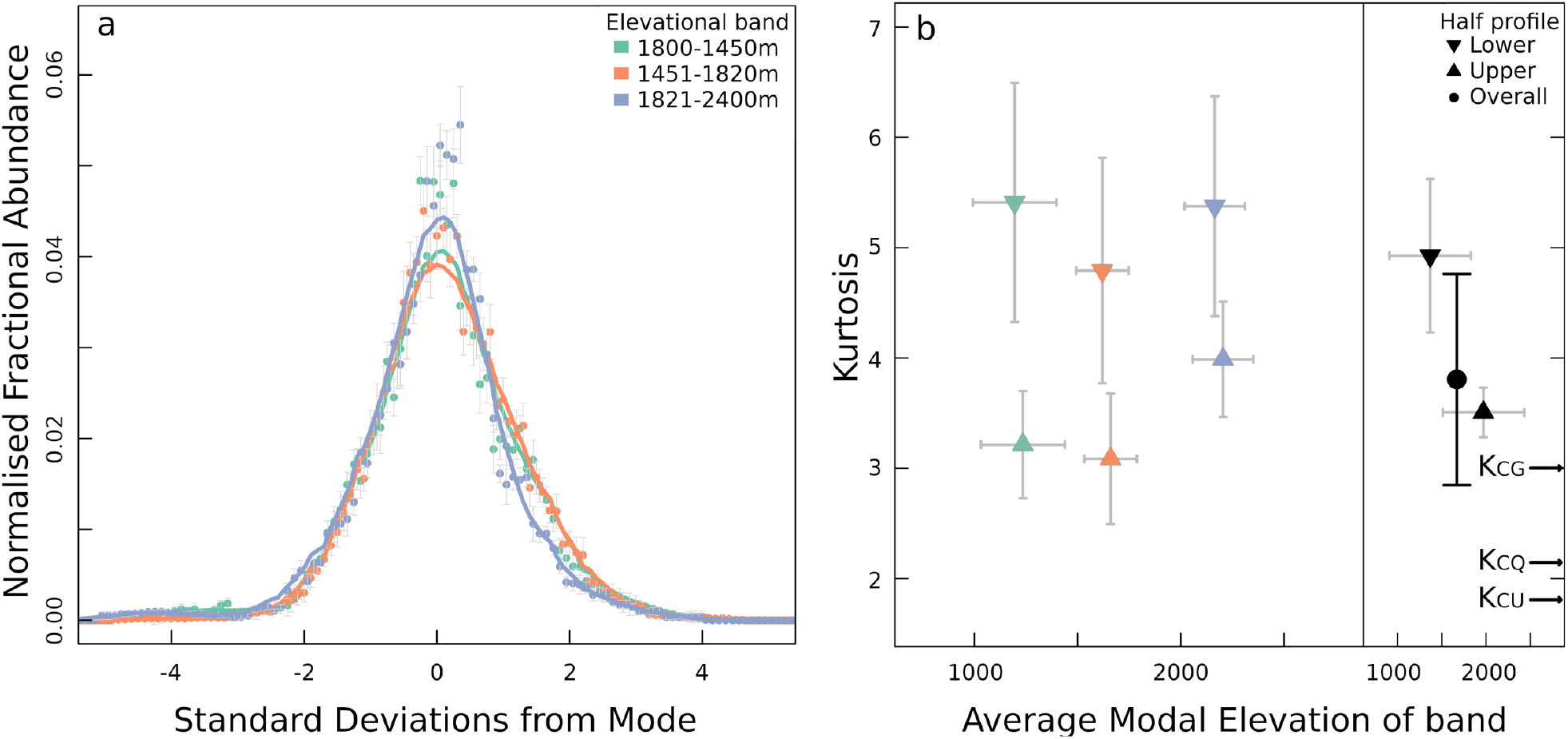
Species averaged community abundance profiles in 3 elevational bands. **(a)** The scatter and smoothed profiles were constructed by averaging the SD-normalised profiles of the species in the community. **(b)** Kurtoses of the half-profiles (split at the mode). The mean and ±1 SE values for each elevational band are shown in colour in the left half of the plot. Their weighted averages and 95% C.I. bars are shown separately for upper and lower halves (3 each) and overall (all 6) in black on the right. The expected smoothed community profile kurtoses are also shown for reference: K_CG_ = 3.0 for gaussian, K_CQ_ = 2.3 for ∩-quadratic, and K_CQ_ = 1.93 for uniform (U_SM_ = 1.93) distributions.

Kurtosis values suggest that the average profile is leptokurtic, i.e. the peak is sharper than for a gaussian, but the tails are heavier. This is also consistent with (i) higher SD for error-normalised A_S_ than for A_60_ (Table 1), and (ii) Figure 6 which shows that a gaussian profile which matches the observed peak is narrower than that which matches the observed SD.

**Figure 6.**
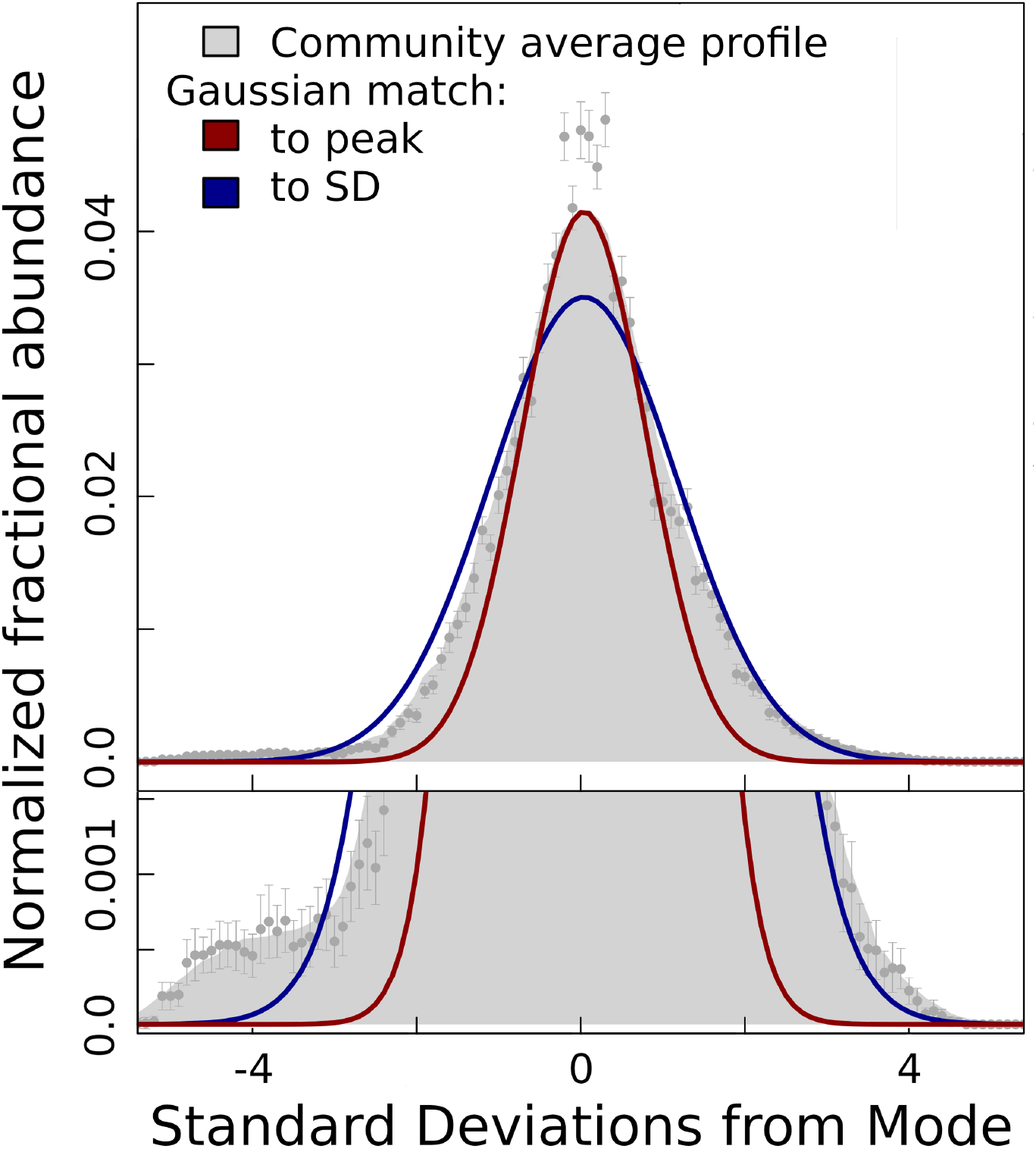
Average abundance profile over all species. SD-normalised profiles of individual species were coadded and smoothed with a width of 1.5 units (grey colour). The red and blue profiles are gaussians matched to the observed peak and SD, respectively. The lower panel shows detail in the outer regions. Only 6 species contribute beyond 3 SD and 1 species beyond 4 SD

## Discussion

We have presented a study of the abundance profiles of bird species in a montane ecosystem in the eastern Himalayas with contiguous primary forest spanning 500-2800 m within a compact region (15 × 6 km^2^ projected area). Departing from the previous heuristic approach we derived ACH as the prediction of a theoretical model applied to an appropriate environmental structure. We recast ACH in terms of the symmetry of a distribution on either side of its abundance peak. We suggest that this is ecologically and methodologically more appropriate than the coincidence of the abundance peak with the geometric centroid of the range defined by the outermost records. We also characterised the range with the more robust root-mean-square-deviation rather than the less reliable width between the outermost records. We found that the mean asymmetry for the community of birds was consistent with zero and the deviaton of individual species profiles from symmetry was consistent with the estimated errors. i.e. the data is consistent with Abundant Centre Hypothesis. There was a negative correlation between the outermost asymmetry metric and elevation. The abundance profiles averaged over species were consistent with a gaussian or leptokurtic profile, while ruling out ∩-quadratic and uniform profiles at a high degree of statistical significance.

### Grafting a Theoretical Framework onto ACH

Despite its insightful and yet simple formulation we have not found any study of ACH in which KB97, or any other model, has been tested with real data. We suspect that this is due to the daunting nature of the differential equations governing spatial distribution of individuals when applied to complex multi-variate environmental patterns in a two-dimensional landscape. Here we tried the alternative strategy of first identifying a “simple” environmental gradient for which KB97 yielded a simplified model and testable predictions. Indeed, ACH emerged as a prediction of the theoretical framework for the particular environmental gradient, of course, under some assumptions (which are discussed below)

#### Single Trait v/s Fitness of the Individual and Multiple Variables

KB97 describes the spatial pattern of distribution of the values of a single trait (i.e. of the individuals with those trait values) across a single-variable environmental gradient. However, the fitness of an organism is influenced by mutiple traits responding to multiple environmental variables. This should not be an issue for several reasons: (i) if the fitness due to a trait results in gaussian abundance profile, it can easily be shown analytically that a combination of traits will also result in gaussian abundance profile. Indeed simulations have shown that *smooth* gradients of mutliple environment variables (and hence multiple response traits) in a 2-dimensional landscape can lead to a smooth unimodal abundance profile (e.g. Brown et al., 1995), and (ii) multiple variables can be reduced to the univariate case if the different variables are strongly correlated to each other.

Multiple environmental factors along an (especially steep) elevational transect are likely to be strongly correlated with elevation. This was certainly true at our study site: a principal component analysis of mean annual temperature, mean annual precipitation, plant productivity, and air density/partial pressure of oxygen yielded a first principal component (PC1) which accounted for 91 % of the variance, and R^2^ = 0.95 for the linear regression of PC1 and elevation (Mungee & Athreya, 2020); i.e. the elevation was an excellent single variable to represent the multi-component environment.

#### One-dimensional Landscape

KB97 describes a one-dimensional profile along a single variable environmental gradient. However, all species ranges are manifestly two-dimensional in geography and respond to multiple environmental variables. A two-dimensional version of KB97 would be much more difficult to solve and of limited utility as an analytical tool. Nevertheless, one can apply one-dimensional analysis (with a suitable change of coordinate system) if the environmental gradient is much lower in the second dimension. This is true of elevational transects, with the environmental gradient very steep perpendicular to an elevation contour and essentially zero along it (e.g. Freeman & Beehler, 2018). For example, in a geographical region spanning just 75 km x 75 km around our study site the elevation changes from 100 m to 5000 m (30°C change in mean temperature) while the highly folded 2000-m elevation contour traverses 1500 km of essentially unchanging environment.

Coastlines have been treated as one-dimensional systems for ACH studies as their length is typically orders of magnitude larger than the width (Sagarin & Gaines, 2002a; Defeo & Cardoso, 2004; Sorte & Hofmann, 2004; Gilman, 2005; Wares & Castañeda, 2005; Samis & Eckert, 2007; Tuya et al., 2008; Rivadeneira et al., 2010; Baldanzi et al., 2013). We suggest that this is inappropriate – transects parallel to the coast are geographically one-dimensional, but not for the purpose of ACH. Coastal ranges of the species in these studies spanned several thousand kilometers with complex variations in multiple abiotic, biotic and anthropogenic factors, and lacked a “unifying” feature like elevation in the mountains. However, in exact analogy with the elevational contours of a montane ecosystem, a transect perpendicular (*and not parallel*) to the coastline is suitable for testing ACH.

#### Symmetry

KB97 linked the symmetry of the environmental gradient to that of the abundance profile – this linkage is at the core of the environment-abundance paradigm. In their model, the trait discrepancy is linear and antisymmetric (not asymmetric; its modulus is symmetric). They explicitly imposed the symmetry by making fitness the square of the discrepancy. In terms of analysis and logistics, the amount of data needed to invalidate a prediction of symmetry is far less than that for falsifying a particular abundance profile. Symmetry can be disproved by showing that some (any) metric is not the same on the two sides of the putative symmetry location (here, the abundance peak). In contrast, testing a predicted profile with data requires sufficient data at multiple locations along the environmental gradient. Therefore, at this early stage of testing theories it would be simplest to identify environmental gradients for which the models predict symmetric abundance profiles. Symmetric abundant profiles may more frequent, or at least easier to identify, in compact one-dimensional and univariate landscapes than in continental-scale, two-dimensional and multivariate landscapes.

#### Trait to Fitness

This is the major hindrance in translating environment and traits to fitness since identifying the environmental optimum for a single trait – let alone multiple traits for dozens of species – is far beyond the scope of present-day knowledge. However, symmetry can mitigate this handicap to some degree. Invoking Occam’s razor we construct a “consistency” argument as follows: a symmetric fitness profile is far more likely to lead to a symmetric abundance profile, than an arbitrary asymmetric fitness profile. Therefore, the detection of symmetric abundance profiles in a transect with symmetric (linear) environmental gradient is more likely to have passed through a symmetric fitness function. This argument is somewhat circular, but self-consistent and the best that can be done in the present day for a quantity (fitness) that can neither be measured directly nor calculated theoretically.

#### Abundance v/s Occupancy and Completeness of Sampling

Sagarin & Gaines, 2002b found that 21 out of 23 separate studies of ACH did not sample the full range of the species investigated. Logistically, this is not surprising: if sampling a range requires N grids along one dimension, it needs N^2^ grids in two dimensions (usually with greater accessiblity issues).

While agreeing that the full range has to be sampled (Santini et al., 2019), we offer a more nuanced and contextual interpretation of “full”. We sampled only a tiny part of a species range in our study (e.g. Figure 1) but we covered its *entire local elevational (hence local environmental) range*. Our objective was not a description of the entire range of environments occupied by the species (which we cannot with this data) but to co-opt theoretical tools to educe quantitative principles of the environment-abundance linkage.

Grid occupancy data from multi-decade surveys such as the North American Breeding Bird Survey or the British Bird Survey (Blackburn et al., 1999; Péron & Altwegg, 2015; Osorio-Olvera et al., 2020) were used to circumvent the resources needed for sampling abundance of wide-ranging species but they are impacted by issues of data heterogeniety and quality (discussed in Santini et al., 2019). We estimated a high dispersion of factor 2-3 in the relationship between occupancy and abundance from a plot in Gaston, 2009. This translates to an uncertainty of 60-78% of the total range in locating the abundance peak for a gaussian profile. Sagarin & Gaines, 2002a have reported differences of up to 50% between published ranges (largely determined by occupancy information) and their own estimates from systematic sampling.

We note even with our large field effort collecting abundance data we only had 44 species for analysis. This is similar to the bird study in New Guinea in which 5000 records yielded only 7 profiles which were completely contained within the sampled range (Freeman & Beehler, 2018).

#### Compact Transect

The 500-2800 m transect in our study fit into a projected area of just 15 × 6 km^2^, all on the southern-most slope of the east-west oriented Himalayas. The temperature difference across this elevational transect corresponds to a north-south (i.e. latitudinal) transect of 2300 km. At these continental scales, many other aspects of zoo-geography, ecological history, regional differences in climate and a patchwork of species-specific “no-go” areas can confound the picture, precluding a simple theoretical model. Our compact site is far less likely to have been influenced by these factors, except for their dependence on elevation, which is our surrogate environmental variable.

#### Appropriate Metrics for Range Widths

The geometric midpoint of the outermost records (hereafter, Min-Max) has no ecological relevance in a non-linear environmental gradient. Vagrants far from the bulk of the population are a regular feature of organisms impacted by ocean and wind currents and human agency. In montane landscapes, an insect can fly, or be blown by wind, across the short distance of its entire range, and beyond, in just one hour. Furthermore, Min-Max data can change considerably with sampling effort and vagrant records. Therefore, any metric referenced to the range edge is likely to be error-prone and may not even be of ecological relevance to the bulk of the species (Gaston, 1990). Suppl. Figure S6 shows results from simulations which attempted to locate the peak of a gaussian (i) by fitting it and (ii) as the midpoint of Min-Max. The fit approach is insensitive to vagrants and becomes more accurate with sample size. On the other hand, the midpoint approach is very sensitive to the fraction of vagrants and does not improve with sample size.

In characterising distribution widths, RMSD scores over Min-Max in several ways: (i) it is defined with respect to a more stable location (abundance peak), (ii) it makes use of the full data set (instead of just two records), (iii) it can be defined even for infinite profiles (e.g. gaussian) which are easier to deal in theoretical models, and most of all (iv) KB97 quantitatively links RMSD of the abundance profile to phenotypic and genetic variance, selection dynamics, heritability, fecundity, intergenerational dispersal and slope of the environmental gradient. Comparing multiple species along the same environmental gradient should help in identifying the role of different traits in determining profile shapes.

#### Environmental Gradient v/s Interspecific Competition

All communities are shaped by a combination of external filters, like environmental factors, and internal filters like competition (e.g. Violle et al., 2012). The former causes a convergence of species traits towards the local community optimum while the latter increases the dispersion of traits within the community. Taking this forward, KB97 describes the impact of the environment without considering species-specific peculiarities and interspecific interaction. Interspecific competition is likely to be a very important determinant of species distribution limits (e.g. Case & Taper, 2000; Price & Kirkpatrick, 2009). Consider two competing species with a zone of overlap (Suppl. Figure S7). Since the impact of any competition is density dependent (Keddy, 1989) one would expect the impact on each species to be higher in the zone of overlap (e.g. Legault et al., 2020). While the precise details of the modification of the original shape may differ from the schematic representation of Figure S3, it is reasonable to assume that the interaction will introduce an asymmetry, but in opposite directions for the two species and with the nett asymmetry for the species pair zero. Therefore, the average asymmetry for the entire community should be a measure of the environmental influence on the shape of profiles. Measuring the individual profiles of just a few species may not correctly reflect the environmental effect.

### Abundance Profiles of the Eastern Himalayan Bird Community in Eaglenest

#### Community Average Abundance Profile

ACH (i.e. A = 0) is the obvious and appropriate null hypothesis for this study. Since a statistical hypothesis cannot be proved, we can at best say that the data is consistent with ACH. Figure 4 and Table 1 suggest that the asymmetry, if any, is small. The average profile of all species (Figure 6) shows a very small departure from symmetry, with a slightly larger gap on one side between the (blue) model gaussian profile and the (grey) data. We recognise that the errors on the asymmetry values of the individual species are large despite the large systematically collected data set. A larger data set may well show a definite departure from symmetry at the level of a few percent.

In any case, as commented by Kirkpatrick while reviewing this manuscript (pers. comm), KB97 used many simplifying assumptions to make the model mathematically tractable. It so happened that we were able to obtain abundance data for birds in a location where the model assumptions were “largely” valid.

The profile of five species showed a large departure from symmetry, well in excess of the formal measurement error estimates. This may be due to interspecific interactions or idiosyncratic life-history traits of particular species. Those aspects are beyond the scope of this paper. We did examine the 15 most abundant species in our data set, but found no congeneric pairs in them (on the assumption that neighbouring congeners are more likely to compete) to test the impact of interspecific competition. It will require a lot more field observations to understand competition networks and larger data sets of those species to address this question.

#### Profile Asymmetry and Elevation

We detected a statistically significant variation of A_S_ (metric using RMSD) with elevation. On the average, species have a larger half-range width on the higher elevation side at low elevations and a larger half-range width on the lower elevation side at higher elevations. i.e. the half-range width is smaller on the side of the nearer elevation limit. We confirmed that this was not an artifact of the sampling limit at 500m and 2800 m by (i) avoiding species with modal elevation below 800 m or above 2400 m, and (ii) investigating any change in asymmetry along the profile in the 5 species where this could have been an issue. The alternative may be that the hard elevation limits (100 m in the Brahmaputra valley, and 3250 m at the ridge) is responsible for this variation.

Despite being in the Shiwalik (Lesser Himalayas) the ridge in Eaglenest is somewhat high at 3250 m. This ridge is akin to a sky island (Warshall, 1995) being 23 km and 40 km away from the nearest 3250 m locations on the main middle-Himalyan range, and isolated from them above the 2275 m contour. It is reasonable to assume that species above 2800 m (nominally) are somewhat isolated. The hard elevational limit is likely to distort abundance profiles by compressing the upper half-range width of the high elevation species, and putting extra pressure on the lower half-range. If this compression cascades downwards through interspecies and intraspecies competition we should expect to see the observed relationship between asymmetry and elevation (Jankowski et al., 2010; Stanton-Geddes et al., 2012; Huntsman & Petty, 2014; Péron & Altwegg, 2015; Wen et al., 2020),. Our highest sampling transects are less than 2 km from the highest ridge and therefore may be close enough to feel the effect. A similar explanation should hold at the lowest elevations as well, because of the abrupt transition over only a few km from lowland hill forests to the tall grass plains of the Brahmaputra valley (Rana et al., 2019). We present this scenario as a point of departure for future investigations; it will require much greater field effort to obtain statistically secure single species profiles and investigate their change with elevation. Alternatively, one can obtain some evidence by comparing species profiles on a mountain with a limiting ridge and another in which the elevation (and habitat) extends well above. Theoretical investigation of the same will require the addition of intra- and inter-specific interaction terms into KB97 (see Case & Taper, 2000; Case et al., 2005; Price & Kirkpatrick, 2009).

Alternatively, the assumption in KB97 that fitness is independent of the sign of the trait discrepancy may not be valid. Higher elevations are thought to be higher stress environments (e.g. Louthan et al., 2015; Cunningham et al., 2016). A trait value which differs from the optimum may have a higher penalty above the abundance peak than below it. This would result in a non-linear trait discrepancy profile. If the curvature of this non-linear function were small it can be replaced by two straight lines of different slopes intersecting at the abundance peak. This naturally leads to our bigaussian model for abundance profiles – each linear “half-gradient” on either side of the peak would result in a half-gaussian whose half-range width is related to its slope. If this explanation were correct the profile asymmetry should be zero at lower elevations and become more negative at higher elevations. However, this is not borne out by our results where the asymmetry is zero close to the midpoint of the elevational range.

We note that a combination of this elevational dependence of asymmetry and the lower number of species above 2000 m (Mungee, 2018) can explain the small amount of asymmetry seen in the average profile (Figure 6).

#### Profile Shape

The kurtosis values of all 6 half-profiles (from averaged profiles in three elevational communities, separately above and below the peak), indicate that abundance profiles have a peak and tails which are at least as broad as that of a gaussian profile. This is in line with the bell-shaped expectation for abundance profiles assumed in many studies (e.g. Hengelveld & Haeck, 1982; Tuya et al., 2008; Boucher-Lalonde et al., 2012; Freeman & Beehler, 2018). Our results clearly reject uniform or ∩-quadratic shapes; i.e. range eges have a tapered profile. We note, again, as pointed out by Kirkpatrick (pers. comm), that this does not “prove” that abundance profiles in nature have to be gaussian. It does demonstrate that theoretical models can reproduce observed data with reasonable assumptions.

Three different approaches showed a “flattening” of the profile from the abundance peak to the periphery: (i) higher SD for error-normalised A_S_, than for A_60_, (ii) variation of A_S_, but not A_60_, with elevation, and (iii) narrower model gaussian when matched to the observed peak than to the observed SD (Figure 6). We recall that A_S_ uses the entire data and has both number and distance, while A_60_ uses the data from close to the abundance peak. This suggests that the small amount of observed asymmetry arises from the small fraction of data at the periphery. We also note that A_N_ does not show an elevational dependence; A_S_ differs from A_N_ in using the distance of a record from the peak. This suggests that the hard ecological elevational limits “push-back” the peripheral populations without appreciably modifying the central regions of the profile and beyond. This may also explain the lack of consensus amongst previous studies, most of which depended on the outermost records to define the distribution centre, and many of which used grid occupancy as a surrogate for abundance.

The heavier tail is unlikely to be due to intraspecific competition since that process would have had a greater impact at the densest part of the distribution; instead the peak is narrower (Figure 6).

Most studies assume that abundance distributions are either uniform or gaussian in shape – the former for the sake of methodological simplicity and the latter because of the ubiquity of the shape in nature. This work shows that abundance profiles are very close to gaussian (Figures 5 and 6), with a small degree of departure at the peak and in the tails.

If the asymmetry arises only from a small fraction of the population does it make any sense to worry about this tail? The symmetric profile in the central parts of the range, encompassing most of the population, may be more relevant for understanding the environment-abundance link; the profile of the periphery is a distraction to be ignored. On the other hand, peripheral populations may be more important for understanding the dynamics of selection and range expansion (Caughley et al., 1988; García et al., 2010; Rehm et al., 2015).

We end the discussion with a comment on KB97. It is obvious that the assumptions used to translate the theoretical framework into testable models were chosen to obtain the desired predictions (e.g. gaussianity and symmetry of profiles). Therefore, this work does not prove that those simple models describe all species profiles. Possibly, other theoretical formulations can be made to yield similar predictions (though we did not find published alternatives) but they are all likely to be variations around the basic theme outlined in KB97. The strength of KB97 lies in its simplicity in putting several ecological processes together in a manner which is mathematically simple and intuitive. This simplicity and the resulting analytical tractability helped us identify an environmental context which may lead to ACH – *this was the key*, which distinguishes this study from previous ones. Should tests of KB97 in other elevational transects prove successful we will have identified a reliable entry-level framework with which to explore abundance patterns in more complex environmental structures, and, result in more refined theories in this field.

In conclusion, ACH is only one of the many features, though perhaps the simplest, characterising the environment-abundance linkage. However, the symmetrical abundance profile implicit in ACH can only arise in environmental gradients with particular characteristics. Theoretical models based on quantifiable ecological processes are essential to identify such ACH-specific environments, and to progress beyond ACH. We suggest that compact elevational transects and transects perpendicular to the coast may be more appropriate for testing ACH. We also suggest that systematic collection of abundance data for a large number of species in such transects may offer the best option for gaining insights into the environment-abundance paradigm.

## Acknowledgements

The field work of this project was carried out with financial support from DST-SERB (Govt of India) and Anand Nadathur. IISER Pune provided diverse financial and logistical support for this research. We thank Deepak Barua for inputs while the analysis was being carried out. We thank Deepak Barua, Sutirth Dey, Umesh Srinivasan, Anand Krishnan, Trevor Price, Robert Colwell, Mark Kirkpatrick and Stuart Pimm for their inputs on a first version of this manuscript. We also acknowledge the generous issue of permits and other support from the Forest Department of Arunachal Pradesh. The field work could not have been undertaken without the help of the Singchung (Bugun) community.

## Contributions

Alakananda Maitra did all the analysis, literature survey and contributed to the manuscript. Rohan Pandit carried out the entire field work. Mansi Mungee helped with the data curation. Ramana Athreya conceived the overall project and supervised the design and execution of the field work and analysis, and led the mauscript writing.

## Supplementary Material for Methods

### Sampling Strategy

All species abundances were recorded by the same observer along 200 m line transects at 47 elevations between 500-2800 m, at equispaced elevational intervals of 50 m (Figure 3; Table S1). Each transect was sampled during a 5+5 minute traverse up and down, on 12 different days between 2^nd^ May and 3^rd^ July, 2012-2014 (Figure S1). All individuals detected (visually and aurally) within 20 m from the path were recorded. Sampling was conducted during 0600-1200 hr, covering up to 12 elevations per day. We minimised systematic bird activity bias by distributing the 12 transects equally across three 2-hour slots – early morning (0600-0800 hr), mid morning (0800-1000 hr) and late morning (1000-1200 hr).

Accessibility issues prevented our sampling along the road below 500 m in Eaglenest. So, we sampled 4 different transects (12 replicates each) at 200 m elevation in neighbouring Pakke Tiger Reserve, which was 25 km away and across the Kameng river gorge. Unfortunately the road in Pakke was laid along an ground contour, unlike in Eaglenest. Given the larger distance to these locations, and the absence of sampling at 250-450 m we only used the 200 m data to identify species whose range extended below 500 m, and exclude them from further analysis.

Having to cover transects spread over ~20 km everyday, the observer used a motorcycle to get one transect to the next. This meant that every transect was traversed twice (labeled, say, A and B) in quick succession to get back to the vehicle. Though we realised that this could result in correlated records we counted birds during the return traverse as well. Furthermore, the probability of sighting the same bird twice varied across the transect, being highest at the far end. We tried two abundance options for each species, viz. A+B ≡ (A+B)/2 and Max(A,B), and could not discern any significant difference in the results apart from the reduction in the number of records. There was no obvious correlation between A and B counts either. So finally, we used A+B as the count for the transect replicate.

One can rationalise this conceptually by recasting our dependent variable: from abundance to the product of abundance and time period of utilisation of the habitat, which will not change any of the final conclusions. We are on firmer ground on the analysis side since correlation between replicates has the same effect on statistical analysis as flocking; it will result in underestimating the counting noise. Since we estimated the counting noise empirically from the data itself (see *Poisson overdispersion factor* below) this partial correlation between the traverses will be reflected in increased noise above the Poisson dispersion.

### Elevational Movement Across The Sampling Period

We examined the data for elevational movement during the sampling period by computing the correlation between ordinal date (regardless of the year) and elevation. Only nine species showed a significant positive correlation in this exercise. Curiously, several species showed a negative trend, which we suspect is a statistical artifact and provides a lower limit on the stochastic nature of the positive correlations. In any case, translating their elevations to a standard date of 30^th^ June made no discernible difference to the results. So we used the uncorrected data in all analyses.

We compared standard deviation (SD), inner 95 percentile width (R_95_) and the more traditional distance between minimum and maximum elevations (R_MM_) as estimators of range width. SD is analytically more tractable while percentile-based estimators are less impacted by outliers. The relationship between the two is well defined for standard functions: e.g. R_95_ corresponds to ±1.96xSD for a normal profile.

### Asymmetry Metrics

The skew is the standard mathematical quantity to estimate asymmetry. However, it depends on the third power of the coordinate and can have large errors. We adopted a split strategy to measure the skew: (i) smooth the observed profile and fit a peak to it. The smoothing makes it easy to fit noisy data (ii) having located the peak we calculated the dispersion separately on either side of the peak (σ_L_ and σ_H_) using unsmoothed data. Our definition of asymmetry is

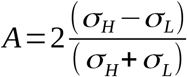

The abundance peak elevation was located using a cubic fit to the smoothed profile. The cubic is the polynomial with the lowest degree which can accommodate (Figure S2). We used the smallest full-width smoothing scale (from among 0.5xSD, 1.0xSD and 1.5xSD) which resulted in a unimodal profile. Profiles which were still multimodal at 1.5xSD were excluded from the analysis.

Our asymmetry metric A is zero for symmetric profiles and ranges between –2 and +2. There are many definitions of skewness in literature – e.g. Pearson’s coefficients (Pearson, 1895), Bowley’s measure (Bowley, 1920). Our asymmetry measure has a simple relationship with prevalent mathematical definitions of skewness (Figure S3). The prior location of the peak using smoothed data and confining our analysis to unimodal distributions, and using the peak location as an input made the skew estimate a bit more secure.

Smoothing will shift the location and height of the peak for an asymmetric distribution. Shifting the peak results in a reduction of the absolute value of the asymmetry, i.e. a bias. We estimated this shift in the peak through simulations. Therefore generated simulated profiles (N = 10000, to minimise stochastic noise) with the following range of input parameters: σ_Li_ : 100-1000 m in steps of 50 m; σ_Hi_ = σ_Li_ -1000 m in steps of 50 m; smoothing width W_S_ = 0.5-3 SD in steps of 0.5 SD. We measured the shift in the peak along with the resulting (ouput) σ_Lo_ and σ_Ho_ and created a look-up-table for reference values of W_S_, σ_Lo_ and σ_Ho_. We used the output values σ_Lo_ and σ_Ho_ for matching since we were unable to calculate the pre-smoothing observed profiles. We used this look-up table to correct for the shift in the observed peak. This smoothing was only used to determine the location of the peak. All subsequent calculations were carried out using the unsmoothed data.

We modeled abundance profiles as bigaussians which is the asymmetric counterpart of the gaussian (Figure S2). For a bigaussian one can estimate the asymmetry A using different parameters all of which provide the same answer (in the absence of noise and non-stochastic outliers). We used

1. root-mean-square-deviation half-widths (analogous to the SD) on the lower (σ_LS_) and upper (σ_HS_) sides of the peak to obtain A_S_. This measure uses all the records in the half-region and each record is weighted by its distance. Therefore, it is most sensitive to distant records.
2. total abundance on the lower (N_L_) and upper (N_H_) sides of the peak to obtain A_60_. This measure uses all the data in the half-region but entirely ignores the distance of the record in its computation.
3. scale lengths at which the smoothed profile falls to 60.65% of the peak value on the lower (σ_L60_) and upper (σ_H60_) sides of the peak to obtain A_60_. The scale length for 60.65% decrement is equivalent to 1 SD in the (bi)gaussian context. This measure is an alternative measure of SD but uses only the ~70% of the data clustering close to the peak.

Estimates of A_S_ and A_N_ are impacted by any section of the profile extending beyond the sampled range. A_60_ is not sensitive to unsampled sections of the profile provided they lie beyond σ_60_. Comparison of A_60_ and A_S_ provide some indication of the change in the relative dispersion of the profile from the peak to the periphery. Comparison of A_N_ and A_S_ identify the contribution of outliers to the observed asymmetry.

One could have used the distances between the peak and the outermost records on the lower (X_L_) and higher sides (X_H_) of the peak to obtain A_MM_. However, A_MM_ will be affected by the same issue that plague the use of X_L_ and X_H_ to determine the centre, and so we did not use this measure

### Poisson Overdispersion Factor

Flocking of birds, weather conditions and habitat heterogeneity may increase the dispersion of abundance counts above the Poissonian. We estimated this dispersion using the difference between the smoothed N_SM_(E) and the observed N_OBS_(E) profiles (Figure S4a). If the observed (raw) profile has an error statistic with standard deviation σ_ε_ in one elevational bin, smoothing it with a window of 5 bins yields an error statistic with standard deviation of σ_ε_ /√5 ≈ 0.447σ_ε_. The difference between the raw and the smoothed abundance is another statistic with mean = zero and dispersion = sqrt(1 + 0.447^2^)σ_ε_ = 1.1σ_ε_; this should be valid for locally linear or low-curvature sections of the profile, i.e. all regions away from the peak. The difference between the errors of the smoothed and the (unknown) “true” profiles is a second order effect and can be ignored. The statistic Y = (N_OBS_ – N_SM_) / √N_SM_ should be approximately standard normal for bins with N_SM_ > 10. However, we estimated σ_Y_ = 2.26 for our data, corresponding to an over-dispersion factor of ~2 (Figure S4b).

### Bird Detectability

The counting error on a Poisson-like process depends on the absolute number of individuals counted in any spatial or temporal interval. In general one is unlikely to spot all the birds of a species within the 20 m strip on either side of a transect. We were only interested in intraspecific comparison of abundance at different elevations. Since all the transects were in a similar habitat structure – along a vehicle track passing through good forest we do not expect much of a variation in detectability of the same species at different elevations. This introduces an unknown multiplicative factor to the actual number of birds in any transect which can change the absolute counts and the error thereon. However, the use of an empirically determined Poisson factor includes the contribution of this unknown detectability factor as well.

### Relationship between Asymmetry and Elevation

Evaluating the statistical significance of the relationship between the dependent (asymmetry) and independent (modal elevation) is complicated because

1. There are errors on both independent and dependent variables.
2. Their errors are correlated. An error in the estimation of the modal elevation will reduce the half-width estimate (N_L_ and N_H_; σ_LS_ and σ_HS_; σ_L60_ and σ_H60_) on the side of the shift and increase it on the other side.
3. The location of the peak is a non-linear function of the empirically-determined profile shape
4. The asymmetry A is a non-linear function of the half-widths
5. The range of A is limited to ±2, because of which its errors will follow some unknown non-standard distribution.

We adopted a layered approach to this question:

1. Determine their non-parametric Spearman’s Rank correlation-coefficient – a very robust parameter which makes no assumptions of the distribution of any input or estimated parameter.
2. Determine the parametric Pearson’s correlation coefficient, whose error estimate is only valid for gaussian errors
3. Determine the linear regression parameters using only the errors of the dependent variable. The fitted slope will be a lower bound of the true value, which works in our favour as it is a conservative estimate of a true relationship.

As it happened, we found a significant relationship from all three tests.

### Profile Shape – Kurtosis

A simplistic recipe to testing if an observed profile is better represented as a gaussian, inverted-quadratic or uniform profile is to (i) fit all three profiles to the observed data by turn, (ii) derive a goodness of fit estimate like, for example, χ^2^ and (iii) use some criteria to pick one of the three as the best fit. This process has two disadvantages which could vitiate the entire exercise: fitting a non-linear curve is not straight-forward and the fitted parameters will end up with large error bars, especially for the quantum of records we have for individual birds.

Instead, we “measured” the shapes using the parameter kurtosis (K) which is characteristic of each family of curves: K = 3.0 for all normal profiles regardless of mean and SD, K= 2.14 for all ∩-quadratic profiles, and K = 1.8 for all uniform profiles.

Kurtosis involves the fourth power of the coordinate in both the numerator and the denominator which results in large errors for small datasets. Therefore, we calculated the kurtosis for species-averaged community profiles in 3 elevational bands (15 species in 800-1450 m, 15 species in 1451-1820 m, and 14 species in 1821-2400 m). The elevational profile of each species was normalised using E_N_ = (E – E_M_) /σ_E_ and F_N_(E_N_) = N(E_N_)/ N_T_, where, E_N_ is the normalised elevation, F_N_ is the fractional abundance at elevation E_N_, E_M_ is the modal elevation, σ_E_ is the elevational SD, N(E_N_) is the unsmoothed abundance at elevation E_N_, and N_T_ is the total abundance for the species. The normalised profiles of all the contributing species (i.e. modal elevation within the band) were averaged at each elevation after weighting it with the inverse of the variance. Finally, the species-averaged profiles were smoothed with full width kernel of 1 unit. We calculated the kurtosis for the 6 half-profiles (above and below the peak) in the 3 elevational bands.

We used simulations to determine the dependence of kurtosis on smoothing width and profile SD (Figure S8). Kurtosis was independent of profile SD for the range of SD values seen in our data (SD: range 60-391 m; mean 241 m).

### Error Estimates

All the parameters we estimated – modal elevation, half-width estimates, asymmetry and kurtosis – depend on the abundance profile in a non-linear manner, and the errors of some of the parameters are correlated. Therefore, we used Monte Carlo simulations to estimate these errors.

We used a model abundance profile, a Poisson overdispersion factor 2.0 (Figure S4) and a negative binomial random number generator (Lindén & Mäntyniemi, 2011; *rnbinom in R*; R Core Team, 2020) to generate 400 simulated profiles for each species.

The model profile for each species was the observed profile smoothed with a width of W_S_ = 5 × 50 m. These simulated profiles were processed in a manner identical to the observed profile to generate 400 sets of modal elevation, the three asymmetry metrics from the corresponding lower and upper half-widths and kurtosis. The simulated distribution function of these parameters were used to identify the error estimates (either standard error or the 95% confidence interval).

## Supplementary – Tables & Figures

**Table ST1.** Mean latitude and longitude of the elevational transects sampled in Eaglenest wildlife sanctuary, Arunachal Pradesh, India. The length of each transect was 200 m. All the transects may also be (approximately) located by matching the altitude on the motorable dirt track (visible on GoogleEarth) in Eaglenest.

| Transect | Latitude (N) | Longitude (E) | Transect | Latitude (N) | Longitude (E) |
| --- | --- | --- | --- | --- | --- |
| 500 m | 27.00701 | 92.42173 | 1650 m | 27.06767 | 92.41605 |
| 550 m | 27.00766 | 92.42071 | 1700 m | 27.06539 | 92.41418 |
| 600 m | 27.00870 | 92.41829 | 1750 m | 27.06664 | 92.41328 |
| 650 m | 27.00533 | 92.41881 | 1800 m | 27.06937 | 92.41210 |
| 700 m | 27.00715 | 92.41587 | 1850 m | 27.06780 | 92.41086 |
| 750 m | 27.00803 | 92.41465 | 1900 m | 27.06750 | 92.40990 |
| 800 m | 27.01668 | 92.41309 | 1950 m | 27.06691 | 92.40400 |
| 850 m | 27.01637 | 92.41285 | 2000 m | 27.07079 | 92.40083 |
| 900 m | 27.01637 | 92.41259 | 2050 m | 27.07447 | 92.40041 |
| 950 m | 27.02145 | 92.41522 | 2100 m | 27.07341 | 92.40132 |
| 1000 m | 27.02133 | 92.41420 | 2150 m | 27.07375 | 92.40589 |
| 1050 m | 27.02269 | 92.41475 | 2200 m | 27.07891 | 92.40399 |
| 1100 m | 27.02287 | 92.41362 | 2250 m | 27.07583 | 92.40607 |
| 1150 m | 27.02394 | 92.41301 | 2300 m | 27.07994 | 92.40588 |
| 1200 m | 27.02448 | 92.41155 | 2350 m | 27.08351 | 92.40527 |
| 1250 m | 27.04585 | 92.41889 | 2400 m | 27.08442 | 92.40474 |
| 1300 m | 27.05472 | 92.41654 | 2450 m | 27.10305 | 92.41693 |
| 1350 m | 27.05953 | 92.41510 | 2500 m | 27.10538 | 92.42126 |
| 1400 m | 27.05903 | 92.41322 | 2550 m | 27.11332 | 92.43560 |
| 1450 m | 27.06009 | 92.41310 | 2600 m | 27.11954 | 92.43487 |
| 1500 m | 27.06066 | 92.41298 | 2650 m | 27.12174 | 92.43560 |
| 1550 m | 27.06164 | 92.41364 | 2700 m | 27.12610 | 92.44663 |
| 1600 m | 27.06304 | 92.41381 | 2750 m | 27.12405 | 92.45195 |
| 1650 m | 27.06767 | 92.41605 | 2800 m | 27.12523 | 92.45511 |

**Figure S1:**
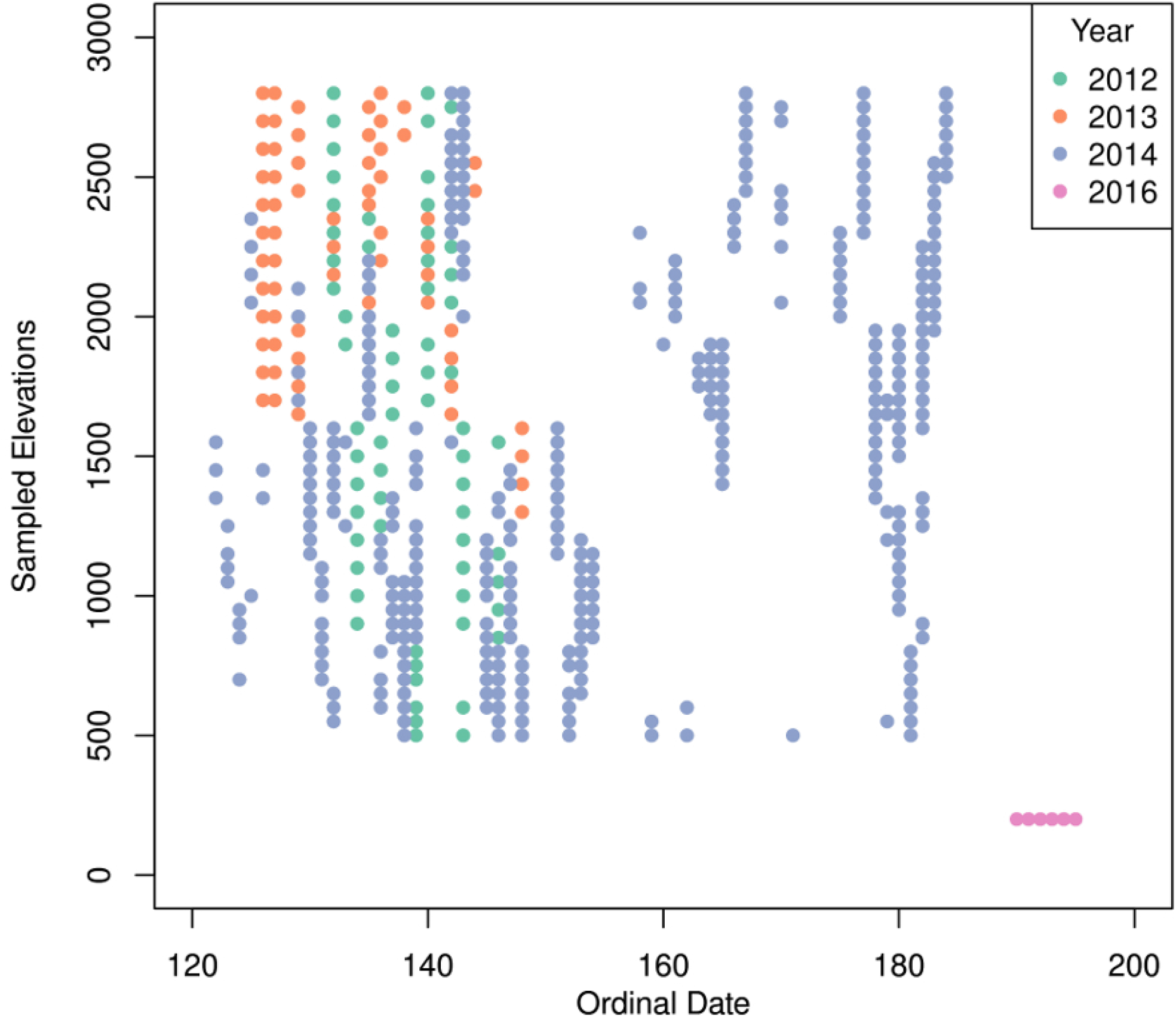
Distribution of sampling dates and transect elevations. The ordinal date values are 122 for May 2^nd^ and 184 for July 3^rd^.

**Figure S2:**
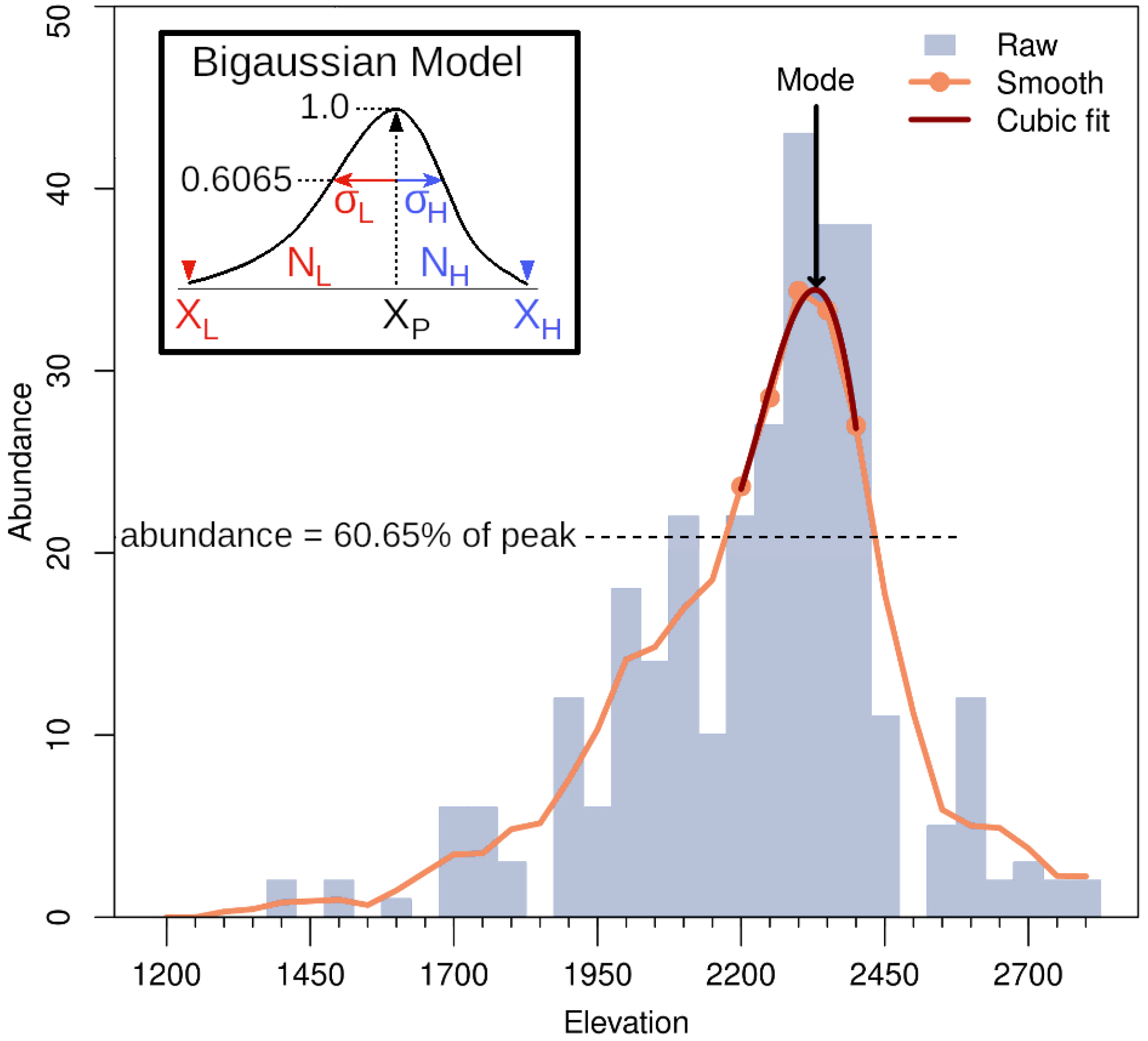
Fitting a peak to the abundance profile. The peak was obtained by iteratively fitting a cubic polynomial to the smoothed abundance profile. At each stage the fit was obtained using elevations with abundance above 60.65% of the smoothed maximum. The inset shows the bigaussian model for the abundance profile. The asymmetry may be defined in terms of the ratio of dispersions (σ_L_ and σ_H_) total numbers (N_L_ and N_H_), or scale length to 60.65% of peak (σ_l60_ and σ_H60_) on either side of the peak.

**Figure S3:**
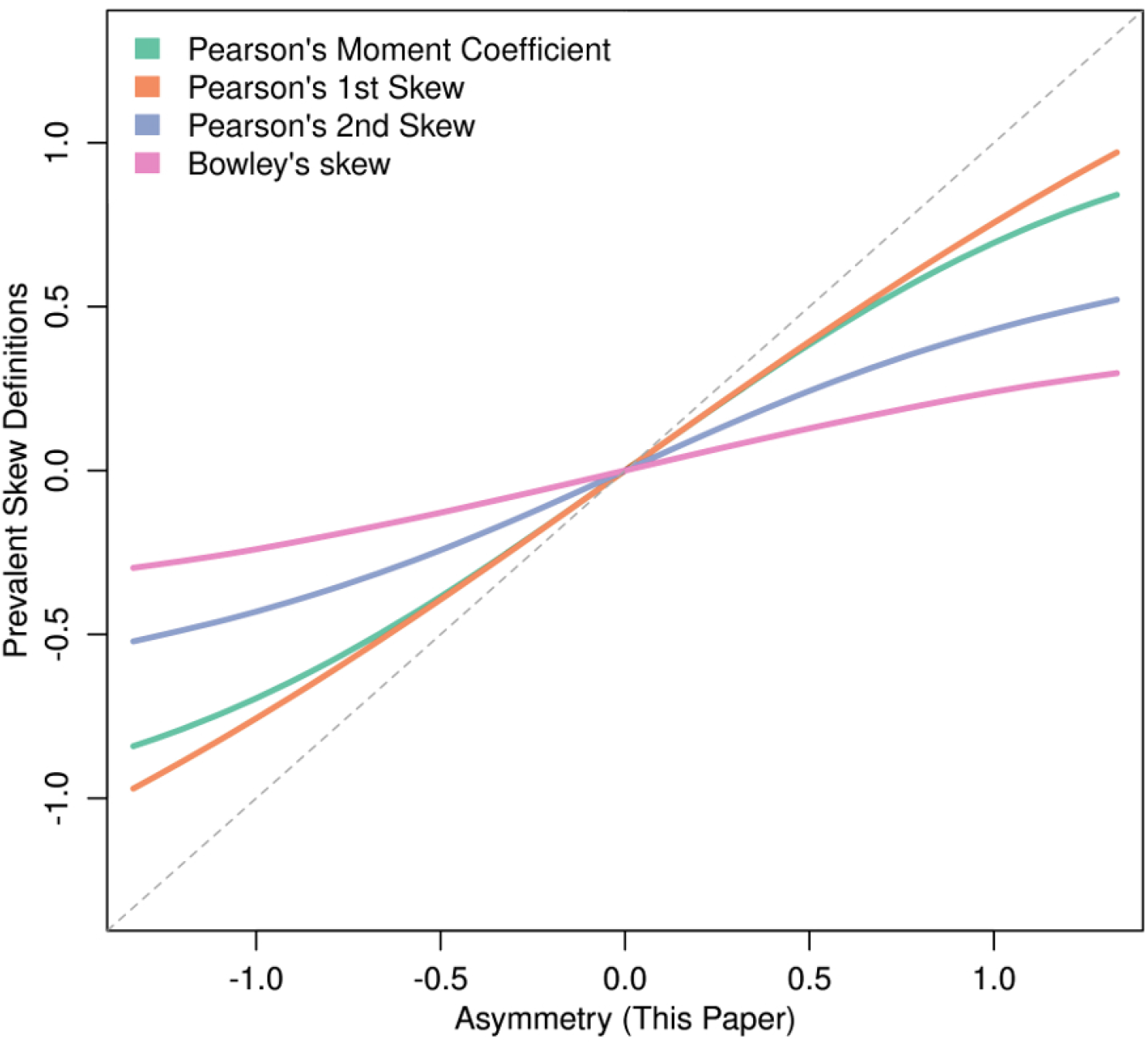
Skewness metrics. Relationship between different mathematical definitions of skewness and the asymmetry metric used in this paper

**Figure S4:**
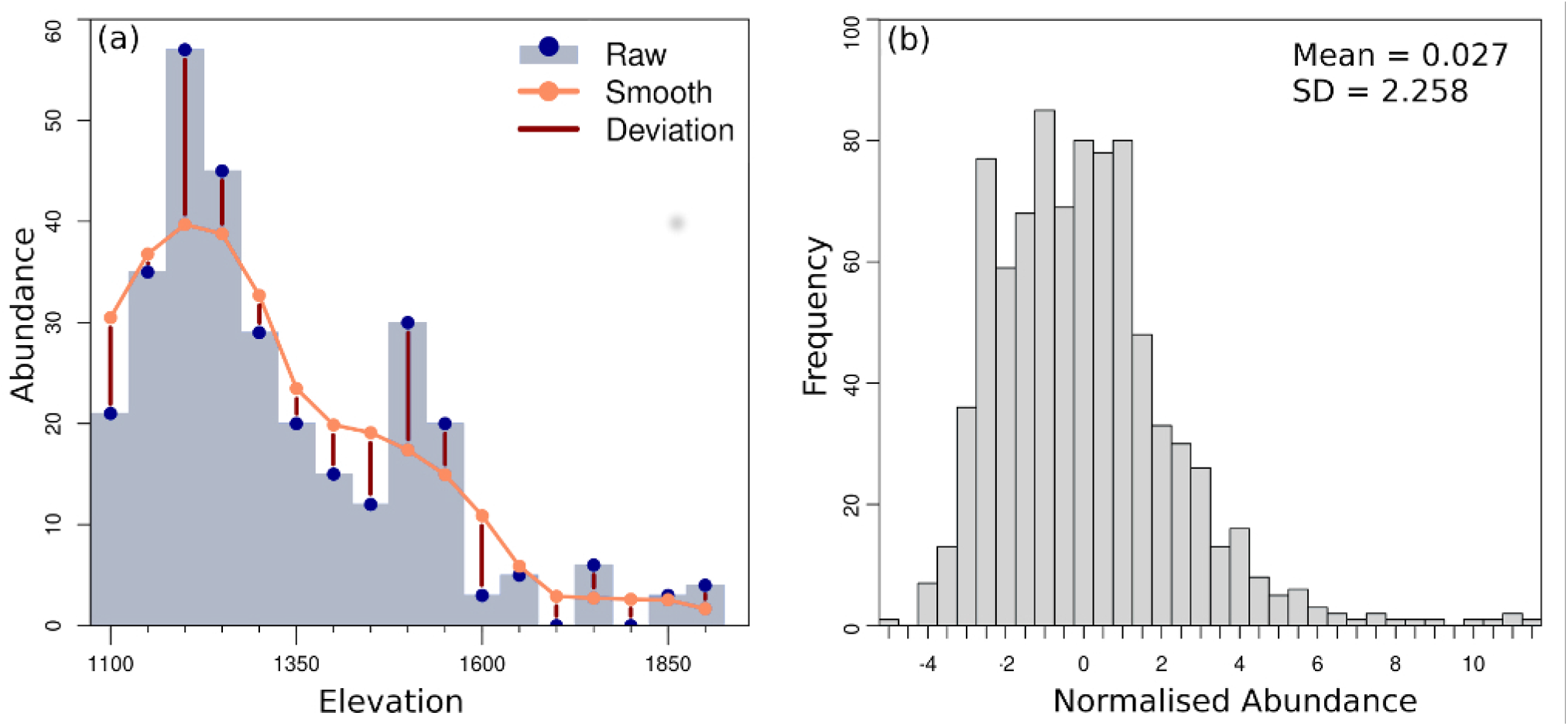
Estimation of Poisson overdispersion factor for abundance. **(a)** Stochasticity in abundance records was estimated using the difference between observed and smoothed abundances in each bin for each species. **(b)** Plot of the Poisson-error normalised deviation statistic. Its mean is close to zero (as expected) but SD = 2.26, which is much larger than that expected for a Poisson distribution (SD=1). From this we estimated that the abundance profiles had a mean overdispersion factor of ~2.0

**Figure S5.**
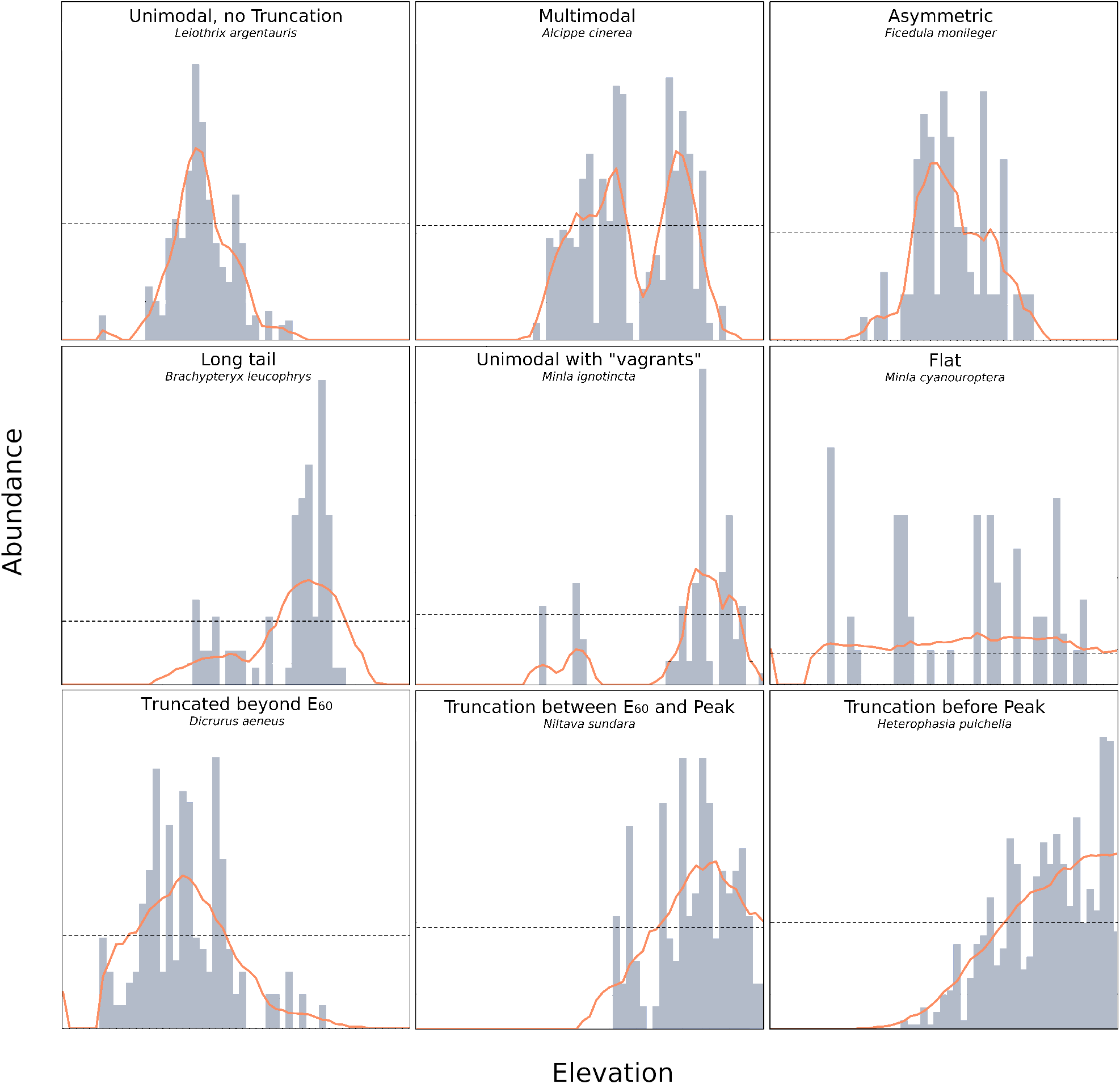
Examples of species abundance profiles in our dataset. The grey bars represent the observed abundances at each elevation, while the orange line represents the smoothed values. The horizontal dotted lines are at 60.65% of peak abundance. Profiles which were unimodal and untruncated were used in the study.

**Figure S6:**
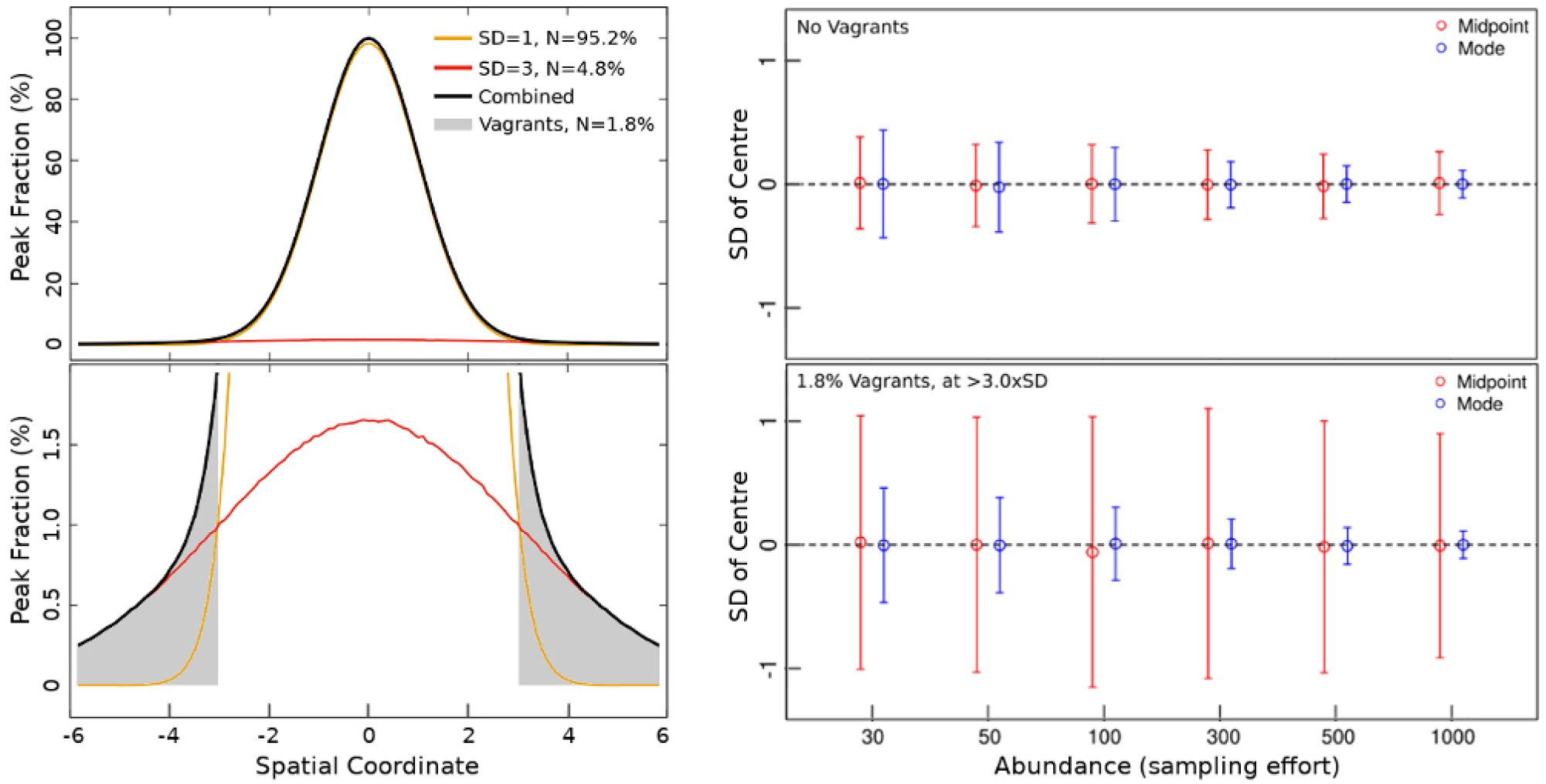
Dispersion in locating the centre of a simulated normal profile. We compared the results by identifying the centre (i) as the midpoint of the outermost records and (ii) by fitting the mode for different abundance values. **Left**: The simulation model (black curve) consists of an admixture of 95.2% “well-behaved” individuals from a normal distribution with SD = 1 (yellow curve) and 4.8% individuals with SD = 3 (red curve). We assumed that all records outside x = ±3 were vagrants (grey-shaded region), since their probability is very low for a “well-behaved” distributions. The ratios were chosen to reproduce the 1.8% “vagrants” in our data. The lower plot shows a magnified section to show the relative distributions of the two components in the outer parts of the range. The composite curve follows the “well-behaved” profile for most of the range and individuals; only a small fraction at the edge makes all the difference. **Right**: The centre coincides with the abundance peak for a normal distribution. The dispersion in the location of the centre from the midpoint of the outermost records (red symbols) is larger than that from mode-fitting (blue symbols), and by a large factor in the presence of vagrants. Increased sampling effort hardly increases the accuracy of the midpoint-as-centre. This is because the centre is estimated from just two records regardless of the sample size.

**Figure S7:**
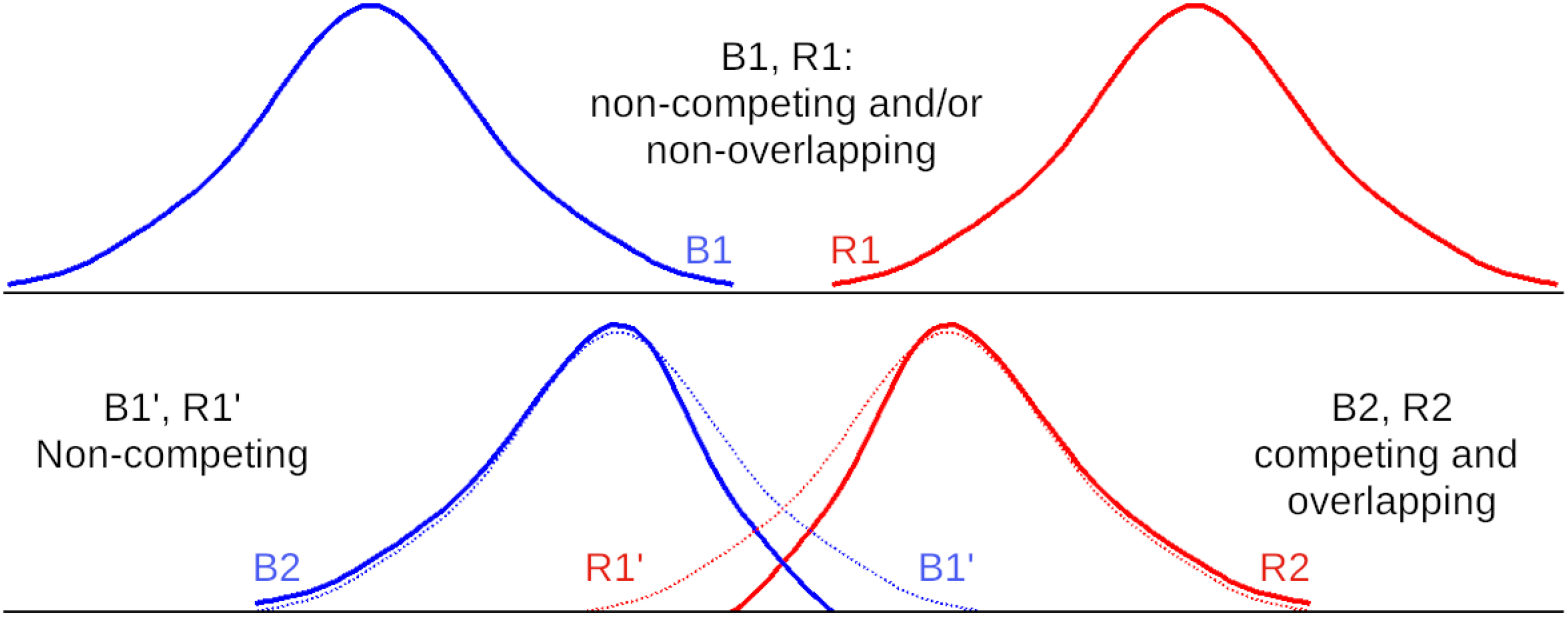
The effect of competition on species abundance profiles. Consider two species for which the environmental component of the shape of the abundance profile is symmetric. Competing species pairs are likely to impact each other much more in the zone of overlap, leading to aymmetric fitness, and hence abundance, profiles. However, the asymmetry will be in opposite directions and the nett average asymmetry introduced by the competitive interaction will be zero.

**Figure S8.**
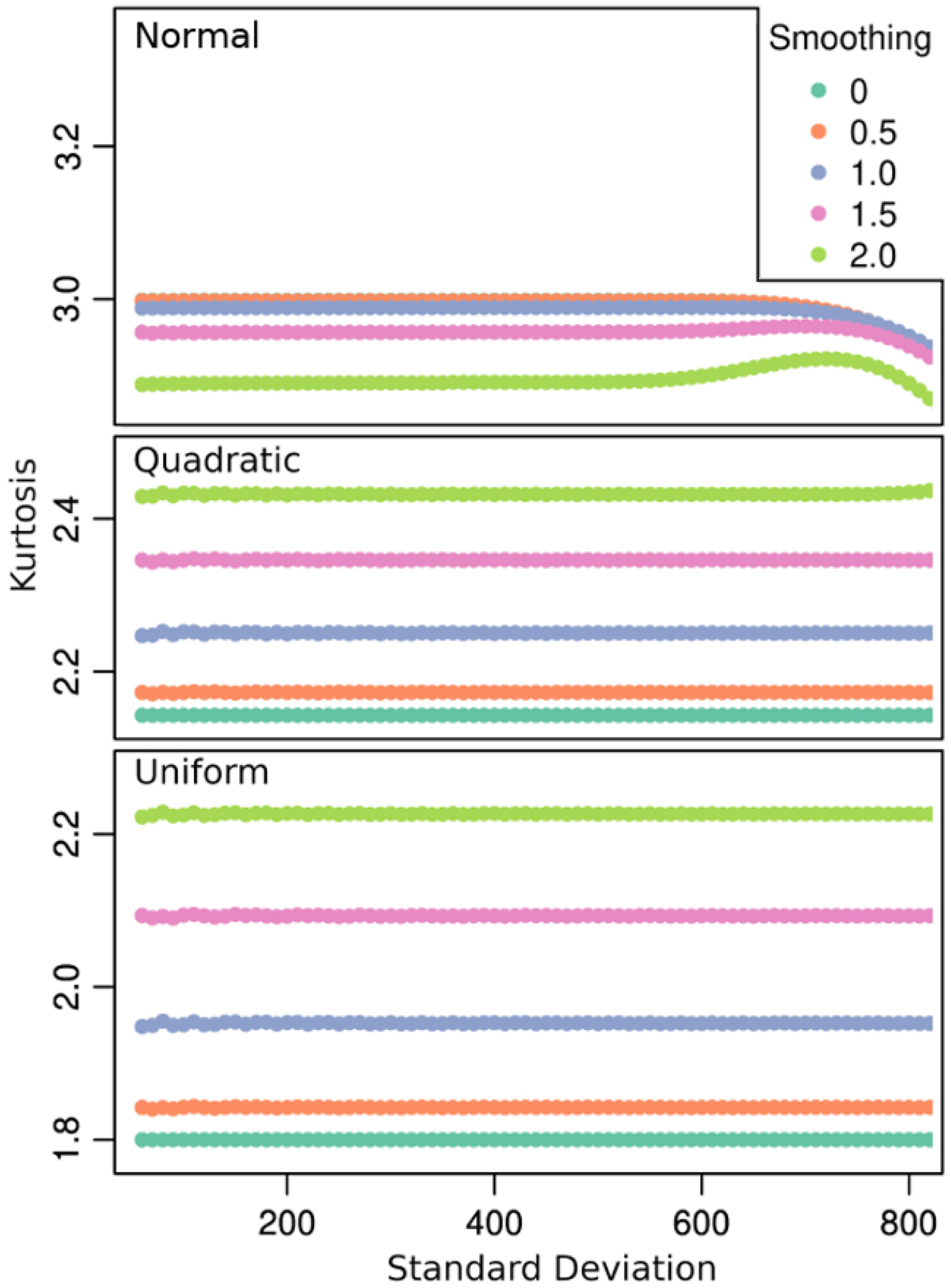
Impact of smoothing on the kurtosis of simulated profiles of different shapes. Colours denote different smoothing full-widths going from 0 (no smoothing) to 2.0 times the standard deviation of the simulated profile. Even with smoothing the kurtosis of nomal (minimum 2.85) and ∩-quadratic (maximum 2.42) profiles are very different. The average predicted kurtosis for our smoothed elevational community samples were normal K_SM_ = 3.0, ∩-quadratic K_SQ_ = 2.23 and uniform K_SU_ = 1.93.

## Footnotes

1 1 We prefer “gaussian” to “normal” to avoid the confusion with the english meaning of the latter

